# CREB-binding protein gene, *HAC701*, negatively regulates WRKY45-dependent immunity in rice

**DOI:** 10.1101/2020.08.26.268797

**Authors:** Nino A. Espinas, Tu Ngoc Le, Miura Saori, Yasuka Shimajiri, Ken Shirasu, Hidetoshi Saze

**Affiliations:** Okinawa Institute of Science and Technology Graduate University, Onna-son, Okinawa, Japan; Bioinformation and DDBJ Center, National Institute of Genetics, Mishima, Shizuoka, Japan; EditForce, Fukuoka, Japan; RIKEN Center for Sustainable Resource Science, Yokohama, Kanagawa, Japan

**Keywords:** *Oryza sativa*, rice CREB-binding protein, *HAC701*, *WRKY45*, innate immunity, *Pseudomonas syringae* pv. *oryzae* (*Pso*), *Magnaporthe*

## Abstract

CREB-binding protein (CBP) is a known transcriptional coactivator and an acetyltransferase that functions in several cellular processes by regulating gene expression. However, how it functions in plant immunity remains unexplored. By characterizing *hac701*, we demonstrate that *HAC701* negatively regulates the immune responses in rice. *hac701* shows enhanced disease resistance against a bacterial pathogen, *Pseudomonas syringae* pv. *oryzae* (*Pso*), which causes bacterial halo blight of rice. Our transcriptomic analysis revealed that rice *WRKY45*, one of the main regulators of rice immunity, is upregulated in *hac701* and possibly conferring the resistance phenotype against *Pso*. The morphological phenotypes of *hac701* single mutants were highly similar to *WRKY45* overexpression transgenic lines reported in previous studies. In addition, we also compared the list of genes in these studies when *WRKY45* is overexpressed and chemically induced transiently with the differentially expressed genes (DEGs) in *hac701*, and found that they largely overlap. When we investigated for *cis*-elements found 1kb upstream of *WRKY45* gene and WRKY45-dependent DEGs, we found that *WRKY45* promoter contains the CRE motif, a possible target of HAC701-mediated regulation. Genome-wide H3K9 acetylation profiling showed depletion of acetylation at large set of genes in *hac701*. However, consistent with the upregulation of *WRKY45* gene expression, our ChIP-sequencing analysis demonstrated that regions of *WRKY45* promoter are enriched in H3K9 acetylation in *hac701* compared to the segregated wild type control in the mock condition. *WRKY45* promoter might be on the receiving end for possible genome-wide compensatory effects when a global regulator like *HAC701* is mutated. Finally, we show that *HAC701* may have roles in systemic immune signaling. We therefore propose that wild type *HAC701* negatively regulates *WRKY45* gene expression, thereby suppressing immune responses.

**SIGNIFICANCE:** HAC701 is a member of CREB-binding protein (CBP) family that acts as transcriptional coactivator and acetyltransferase. However, little is known how it regulates innate immunity in plants. Herein we reported that rice *HAC701* suppresses WRKY45-dependent defense pathway. Our study showed that *HAC701* seemingly interacts genetically with *WRKY45* in rice to modulate immune responses against pathogens.

## INTRODUCTION

Histone acetyltransferases (HATs) are a diverse group of histone- and non-histone-modifying enzymes that contain multiple subunits to enable catalytic functions (1). These HATs primarily function as scaffolding bridges of proteins to form the transcriptional regulatory complex important for target recognition and subsequent substrate modification (1–3). The association of HATs to its interacting proteins and the existing state of the epigenome dictates the dual-switch functionality of either transcriptional activation or repression (1). They are considered integrators or adaptors as they were shown to interact with DNA-binding activators and the basal transcriptional machinery (4, 5).

In plant model Arabidopsis, HATs and histone deacetylases (HDACs) are grouped into four (Figure 1A) and three families, respectively (6). HATs consists of CBP, TAF1/TAFII250, GNAT, and MYST, while HDACs are RPD3/HDA1, HD2-like, and SIR2. A previous review of HATs and HDACs in plants listed about 12 putative HAT and 18 HDAC proteins exhibiting acetylation and deacetylation activities with differential site specificities (6). Work on rice (*Oryza sativa*), however, has just begun with a preliminary investigation of eight histone acetyltransferases (7, 8) and deacetylases (9).

**Figure 1.**
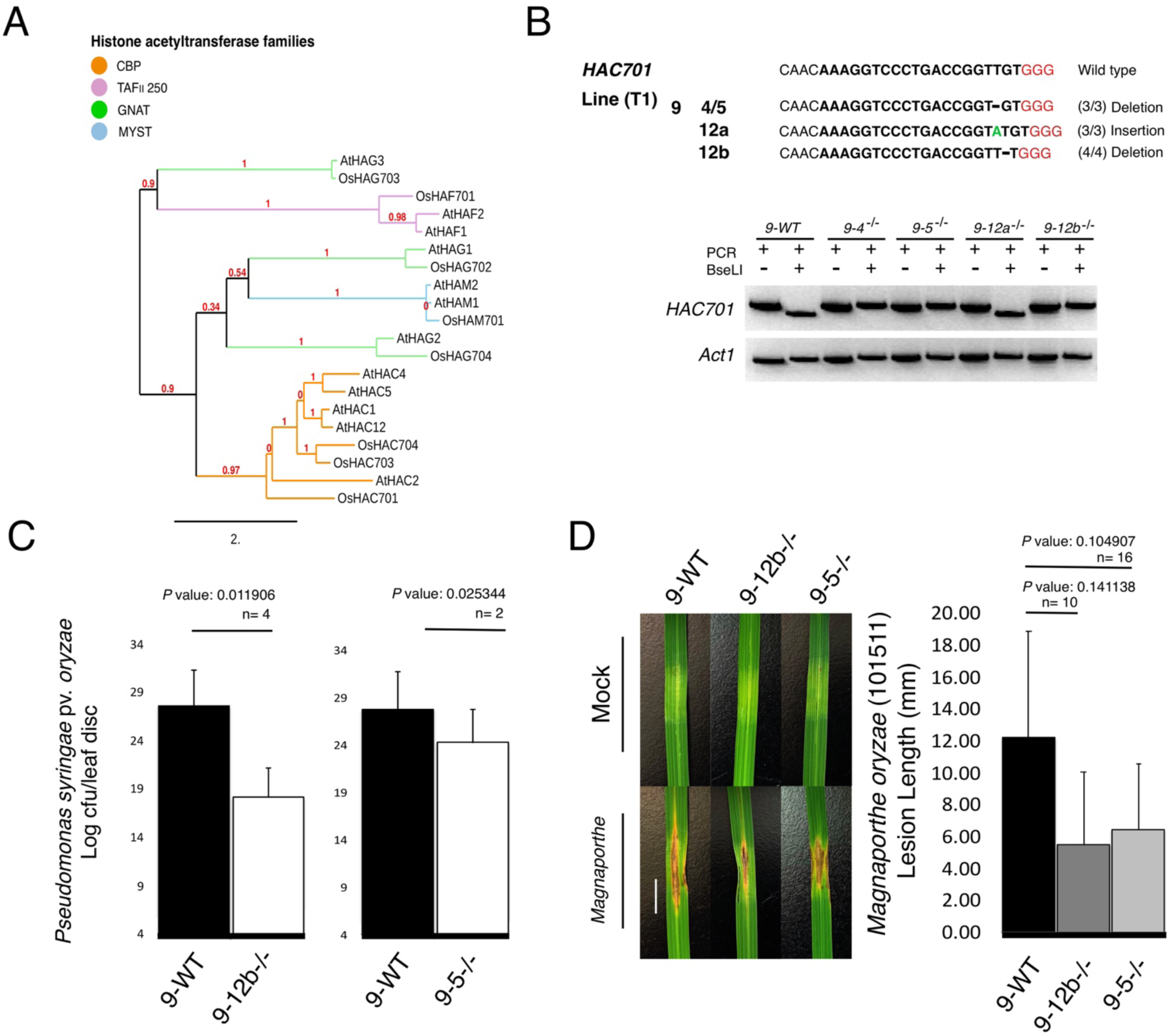
Disease phenotype of biallelic homozygous CRISPR/Cas9 lines targeting the fifth exon of rice acetyltransferase gene, *HAC701*. (*A*) Phylogenetic relationships of members of Arabidopsis and rice histone acetyltransferases. The numbers in red are branch support values. All amino acid sequences are from UniProt database. (*B*) Upper panel: Alleles from four lines of the second generation (T1) plants were identified by cloning and sequencing the PCR products from *HAC701* target region using the primers found in Table S1. Lower panel: PCR and RFLP assays of representative T1 generation lines. The mutant line *9-12a^-/-^* displaying a wild type-like band was due to unaltered sequence order recognized by BseLI even after an insertion event, however we confirmed by sequencing that an insertion mutation was present in this line. (*C*) *Pseudomonas syringae* pv. *oryzae* (*Pso*) infection assay of *hac701*, *9-12b^-/-^* and *9-5^-/-^* (T2), using *9-5^+/+^* segregated wild type line (9-WT) as control. Bacterial density quantification uses values from 4^th^-6^th^ serial dilutions. The mutant line *9-12b^-/-^* was grown from the seeds of the third generation (T2), while *9-5^-/-^* line was embryonically rescued about 15 days post flowering from the second generation (T1) parental plants. All mock measurements yielded zero bacterial growth for both wild type and mutant lines. Bars represent the 95% confidence interval (CI) and compared to the wild type using two-tailed Student’s *t*-test at *P*<0.05 in *9-12b^-/-^*; one-tailed Student’s *t*-test at *P*<0.05 in *9-5^-/-^*. (*D*) *Magnaporthe oryzae* infection assay of *hac701*, *9-12b^-/-^* and *9-5^-/-^*, using *9-5^+/+^* segregated wild type line (9-WT) as control. Lesion length was measured on leaves after 10 dpi (days post infection). White length indicator bar on the left panel is 10 mm. Bars on the right panel represent standard error (SE) and compared to the wild type using *F*-test for variance and Student’s *t*-test.

Several studies have shown that histone acetylation and deacetylation control defense signaling in plants in response to phytohormone or pathogen application (10–12). More specifically, HATs and HDACs in rice model were shown to be responsive to abiotic stresses and can be modulated by phytohormones, thus implicating a role in biotic stresses (7, 13). Similar to Arabidopsis, HATs in rice are also classified into four families (Figure 1A) (7). In contrary, rice HDACs are represented in only two families, RPD3/HDA1-like and SIR2-like with no known rice member belonging to HD2 family (9).

*OsHAC701* was reported (14) earlier as a putative rice CBP-related acetyltransferase, herein after referred to as *HAC701*, of the p300/CBP acetyltransferase (PCAT) family of proteins. The Arabidopsis genome contains five CBP genes, while rice genome contains three (Figure 1A), having broad acetyltransferase specificity on histones (7). Currently, there is no clear consensus on the similarity of biological functions of Arabidopsis and rice homologs of CBP family proteins. In animals, p300 and CBP are paralogs and originally described as transcriptional coactivator that exhibit histone acetyltransferase activity on all four core histones specifically at H4 N-terminal tail sites K5, K8, K12, and K16 (15, 16). Other sites include H3 sites K14, 23 (17), K18, K27 (18, 19), and K56 (20); H2B sites are K12 and K15 (21). H3K9 is mainly acetylated by GCN5/PCAF (18), however acetylation by p300/CBP was also reported (17, 22, 23). Its highly conserved function is mainly found in multicellular organisms as it probably participates in complex physiological processes acting as a limiting factor in various pathways due to its high cellular demand (14).

As HATs have been shown to be modulated by biotic stresses (24), we hypothesized that it could be a potential system for studying rice immunity. Indeed, we show that *HAC701* is involved in rice innate immunity. In this study, we have made CRISPR/Cas9 edited *hac701* lines and showed that *HAC701* is apparently a negative regulator of immunity with the mutants showing resistance against bacterial and fungal infection. Further transcriptomic investigation showed that HAC701 possibly regulates *WRKY45* loci, one of the main regulators of rice immunity pathways. In *hac701*, *WRKY45* and its direct and indirect gene targets were upregulated upon pathogen inoculation. In addition, *hac701* phenocopies *WRKY45*- overexpression (*WRKY45-ox*) lines, indicating a potential interaction between *HAC701* and *WRKY45*-dependent immunity pathway in rice.

## RESULTS

### Characterization of a rice acetyltransferase mutant, *HAC701*

We isolated null mutant lines by independently targeting two sites of *HAC701* gene (Fig. S1A-C; Fig. S1D-I) using CRISPR/Cas9 editing technology. From the 11 positive *HAC701* lines (Target 2), we have isolated four biallelic homozygous mutant lines (Fig. S1D), which we utilized for experimentation including their segregating wild type siblings, 9-WT (Fig. 1B). The phenotype in terms of effective grain number and tiller number of mostly all biallelic homozygous mutants at T2 generation did not show impairment as compared to 9-WT (Fig. S1F). In addition, the T3 lines did not show any visible growth or developmental defect in height and photosynthesis (*e.g.* chlorosis phenotype) (Fig. S1G), respectively. However, the weights of 1000-grain T3 and T4 generations of mutants were increased compared to 9-WT (Fig. S1I). These results indicate that a loss of HAC701 does not cause severe developmental defects in rice, and that HAC701 may have a positive role in seed production.

### *HAC701* mutant rice is resistant to pathogen infection

Our expression analysis of the rice HAT genes showed that *HAC701* is significantly up-regulated by treatment of bacterial flg22 peptide, suggesting a potential role of HAC701 in plant defense against pathogens (Fig. S2). To investigate the involvement of *HAC701* in basal defense in rice, we inoculated 9-WT and *HAC701* mutant lines, *9-5^-/-^*, *9-12a^-/-^*, and *9-12b^-/-^* with mock (10 mM MgCl_2_) and *Pseudomonas syringae* pv. *oryzae* (*Pso*) (OD = 0.2; resuspended in 10 mM MgCl_2_) for 72 h. *Pso* is a causative agent of halo blight in rice characterized by brown lesions and yellow halo-like blotches on leaves (25). Infection of *Pso* showed that *hac701* was resistant to *Pso*-treatment as compared to 9-WT (Fig. 1C; Fig. S3). These data suggest that *HAC701* might be involved in the regulation of rice innate immunity acting in its capacity as a transcriptional coactivator and/or as an acetyltransferase. We further tested whether *hac701* also shows resistance phenotype when infected with a rice blast pathogen, *Magnaporthe oryzae*. Although the data were not statistically significant, results showed a resistant phenotype tendency in *hac701* (Fig. 1D). Overall, these indicate that *HAC701* plays a negative role in rice innate immunity.

### Transcriptome profiling of the rice-*Pso* pathosystem

To profile the genome-wide effect of *HAC701* mutation in rice innate immunity, we performed RNA-sequencing on mock- and *Pso*-treated 9-WT and *hac701* using local leaf tissues (Fig. S4; Table S2). We first characterized the effect of *Pso* infection on 9-WT and analysis of the highly variable genes in mock- and *Pso*-treated 9-WT plants showed gene clusters that were dependent on *Pso* induction alone (Fig. S4; Fig. S5). The top 10 highly variable genes across the 9-WT samples showed features involved in tolerance and/or resistance such as disease resistance and stress tolerance mostly by catalysis of primary and secondary metabolism (Fig. S5). This includes well-documented genes involved in plant resistance against pathogens, such as tryptophan decarboxylase 1 (Os08g0140300) that catalyzes the production of serotonin via conversion of tryptophan to tryptamine in rice (26), which has been implicated to confer resistance in rice infected with *Bipolaris oryzae* (27). Another gene, naringenin 7-O-methyltransferase (Os12g0240900), that catalyzes the production of rice phytoalexin sakuranetin has been reported to participate in rice defenses via JA signaling (28, 29). Lignin and phytoalexin encoding genes for instance laccases (*e.g.* Os12g0258700, Os11g0641500) and cytochrome P450s (*e.g.* Os07g0218700, Os08g0508000) are metabolic products known to be involved in plant defenses as well (30–32). Gene Ontology (GO) analysis of upregulated genes in *Pso*-infected 9-WT plants revealed enrichment of genes involved in response to biotic stimulus (GO terms: Response to wounding, Response to other organism, Cell wall macromolecule catabolic process, *etc.*) (Fig. S6; Table S3). Both up and downregulated genes showed enrichment of transcriptional machinery regulation, a possible consequence of defense-related transcriptional reprogramming processes during bacterial invasion. GO analysis also identified pathways associated with upregulated gene networks involved in plant defense such as diterpene phytoalexin biosynthetic pathway and chorismate biosynthesis. Thus, the rice-*Pso* pathosystem is a functional system to analyze defense responses of rice against *Pso* infection.

### Potentiated *WRKY45* gene in *hac701* provides resistance against *Pseudomonas* and *Magnaporthe* infections

To investigate the molecular basis of resistance phenotype observed in *Pso*-infected *hac701*, we further analyzed our transcriptome data and found 141 upregulated genes in *hac701* (Fig. S7A; Table S4). We refer to these genes as *HAC701*-repressed genes. GO analysis of *HAC701*- repressed genes showed enrichment of processes involved in response to xenobiotic stimuli and defense response (GO terms: Response to xenobiotic stimulus, Response to wounding, Regulation of defense response, Jasmonic acid mediated signaling pathway, Defense response, *etc.*). These genes were lowly expressed in the 9-WT background. On the other hand, 336 upregulated genes in the 9-WT background alone referred to as *HAC701*-independent genes did not show a very distinct enrichment of genes in defense responses yet rather showed a more general biological and physiological processes (GO terms: Mitochondrial respiratory chain complex IV assembly, Alanine transport, Cellular response to heat, *etc.*) (Fig. S7A; Table S3). We also looked into the features of 31 downregulated genes in the mutant background referred to as *HAC701*-enhanced genes (Fig. S7B; Table S4). These genes were not normally downregulated in the 9-WT background. *HAC701*-enhanced genes contained genes enriched in photosynthetic and response to abiotic stress terms (GO terms: Glycerolipid biosynthetic pathway, Response to cold, Photosynthesis, *etc.*). In addition, the remaining 46 downregulated genes in the 9-WT background alone also showed a more general biological processes with no hint of enrichment in defense response against pathogen infection (GO terms: Chloroplast organization, Gluconeogenesis, Organelle fission, *etc.*) (Fig. S7B; Table S3).

We were intrigued what genes were affected in *HAC701* mutant background without *Pso*-treatment. Thus, we analyzed the transcriptome of mock-treated *hac701*, and found a small number of DEGs (98 genes; Table S5) mainly enriched in NADH dehydrogenase complex assembly and photosynthesis. This is in contrast with the number of DEGs found in *Pso*-treated *hac701* transcriptome, which yielded 660 genes (Table S6). These results simply indicate that *Pso* induced the expression of substantial number of genes in *hac701*. To fully understand the role of *HAC701* in *Pso*-treated conditions, we overlapped the list of genes derived from mock and *Pso*-treated mutants, and found *WRKY45* (*OsWRKY45*) upregulated in the *hac701* background in mock-treated *hac701* (Fig. 2A). *WRKY45* was also identified as one of the 141 upregulated genes in *Pso*-infected *HAC701* mutant (Fig. S7A). There was no candidate gene found downregulated in *HAC701* mutant background alone (Fig. 2A). We then checked whether there were differences in the upregulation strength of *WRKY45* in the *HAC701* mutation alone, compared to *HAC701* mutation combined with *Pso*-treatment. Transcriptome data revealed that *WRKY45* expression was increased by about 20% upon *Pso*-treatment (Fig. 2B). These results suggest that *WRKY45* expression is partially dependent on *HAC701* mutation. In addition, *Pso*-treatment allowed the co-expression of genes downstream of *WRKY45* pathway in a HAC701-dependent manner.

**Figure 2.**
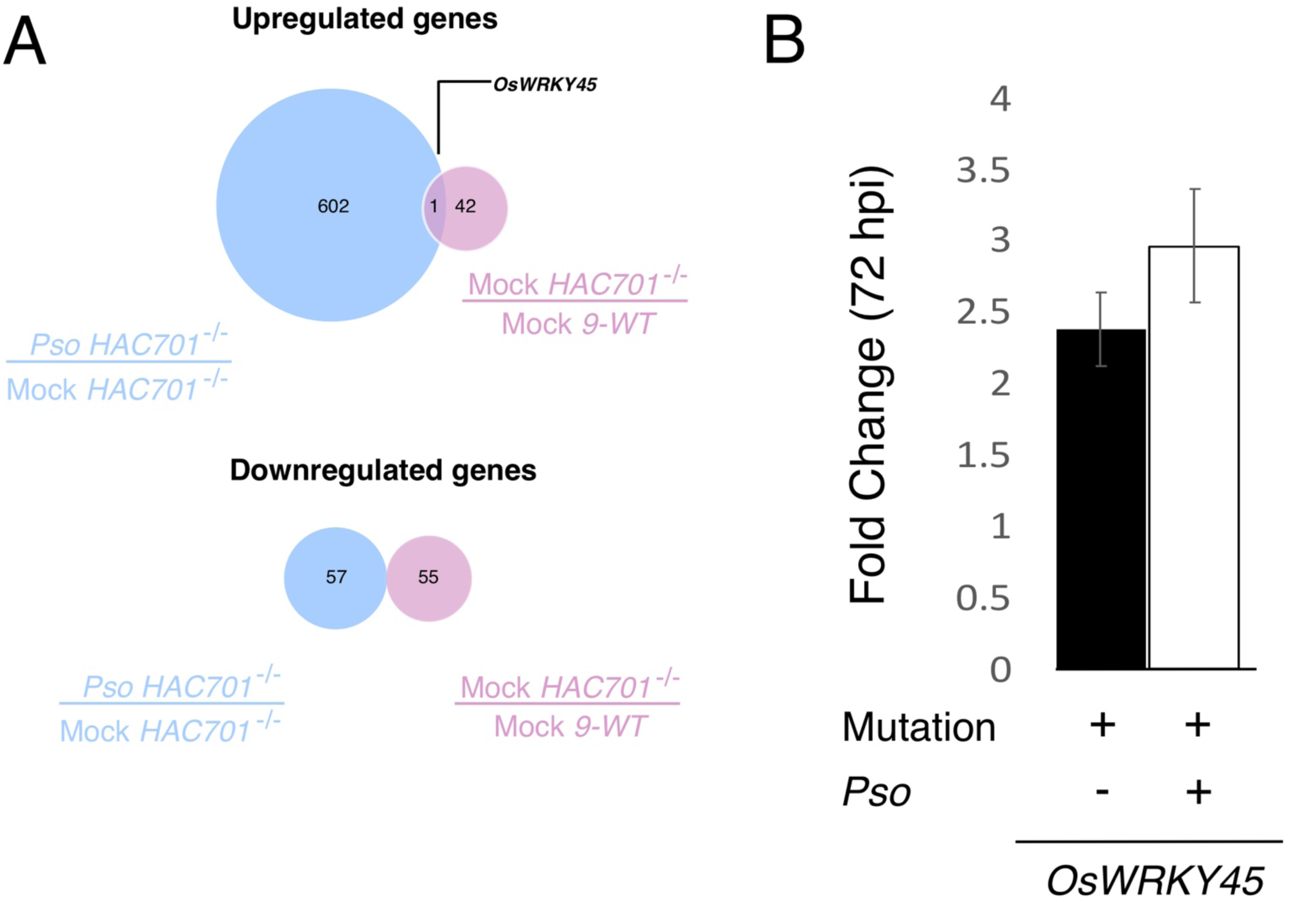
The rice *WRK45* gene expression is induced in *hac701* lines. (*A*) Overlap analysis of upregulated and downregulated genes in *hac701* background under mock and pathogen (*Pso*)- treatment conditions (Table S5, S6). *OsWRKY45* is the only differentially expressed gene in either the absence or presence of pathogen. Transcriptome data were normalized using mock data sets for each genotype and condition. (*B*) *OsWRKY45* expression when *HAC701* is mutated (+ sign). The addition of pathogen (+ sign) on *hac701* increased further the *OsWRKY45* expression. Transcriptome data of *hac701*, *9-12b^-/-^* and *9-5^-/-^*, under mock or *Pso*- treated conditions were lumped as two independent biological samples for analysis (See Table S2). hpi, hours after infection.

### *hac701* phenocopy *WRKY45* overexpression transgenic plants

To determine whether the upregulation of *WRKY45* in *hac701* is relevant to rice with overexpressed *WRKY45*, we compared the DEGs in our *hac701* with *WRKY45* overexpression (*WRKY45-ox*) lines previously reported (in these studies: 33, 34). Our results indicated a striking similarity of targeted genes from rice samples despite the fact that they were inoculated with two different types of pathogens, a bacterial (*Pso*; this project) and the fungal pathogen *Magnaporthe* (33, 34) and minor differences in sampling time periods (Fig. 3). In addition, we then analyzed whether benzothiadiazole-responsive genes (Fig. 3A; Table S6; Table S7) would overlap with *HAC701* mutant-DEGs in *Pso*. It is known that benzothiadiazole (*i.e.* BTH is a synthetic analog of salicylic acid, SA) robustly induces defense genes in rice (34, 35), which usually contains high levels of SA. As a result, about 43.3% (286/660 genes; *P*<3.336e-161) of *HAC701* mutant-DEGs in *Pso* overlapped with BTH-responsive genes, while about 12.4% (82/660 genes; *P*<2.368e-79) of the *HAC701* mutant-DEGs in *Pso* overlapped with BTH- and WRKY45-regulated genes (Fig. 3B; Table S8). Furthermore, genes upregulated in plants expressing DEX-induced myc-tagged WRKY45 protein in rice were mostly found in *HAC701* mutant-DEGs in *Pso* (75% or 9/12 genes; Table S9). Our analysis also showed that at most 9.1 % (60/660 genes; *P*<3.986e-15) of the *HAC701* mutant-DEGs in *Pso* were regulated in BTH- and NPR1/NH1-dependent manner (Fig. 3B; Table S10). The rice NPR1/NH1 is an independent immune signaling pathway that mediates gene responses through BTH/SA-induction (Fig. S18). We then analyzed the 72 *HAC701* mutant-DEGs in *Pso* that were exclusively regulated by BTH- and WRKY45 as well as the other 10 genes jointly regulated by WRKY45 and NPR1/NH1 under BTH treatment (Fig. 3B-D). We found that most of these *HAC701* mutant genes were genuine WRKY45-regulated genes (Fig. 3C-D) (33, 34). Analysis of these WRKY45-dependent defense-related genes showed that in the presence of *Pso*-infection, a number of glutathione-S-transferase (GST) gene, were more upregulated in 9- WT than in *hac701* (Fig. 3D). These results were also consistent with the result that *OsWRKY62*, a known negative regulator and direct target of *OsWRKY45* (34), had lower transcription level in *HAC701* mutants (Fig. 3D). It is interesting that genes in JA biosynthesis, signaling and perception tend to be more highly expressed in *HAC701* mutant than 9-WT (Fig. 3D). In addition, *HAC701* mutant genes that were found regulated in BTH and *WRKY45* indeed contain W-box motifs (i.e. binding motifs for WRKY transcription factors, 36), raising more the possibility that these genes are direct WRKY45 targets (72/82 or 87.8%; Table S11). Meanwhile, all 10 genes that were commonly regulated in BTH, *WRKY45*, and *NPR1/NH1* (Fig. 3) contain W-box motifs as well (10/10 or 100%; Table S12). Taken together, these results indicate that upregulation of *WRKY45* expression as an outcome of *HAC701* mutation resulted in expression profile of genes almost identical to overexpressing *WRKY45* transgenic lines as well as to DEX-inducible *WRKY45* expression system. These also suggest that a significant fraction of *HAC701* mutant DEGs are probably WRKY45-regulated, given the presence of W-box motif.

**Figure 3.**
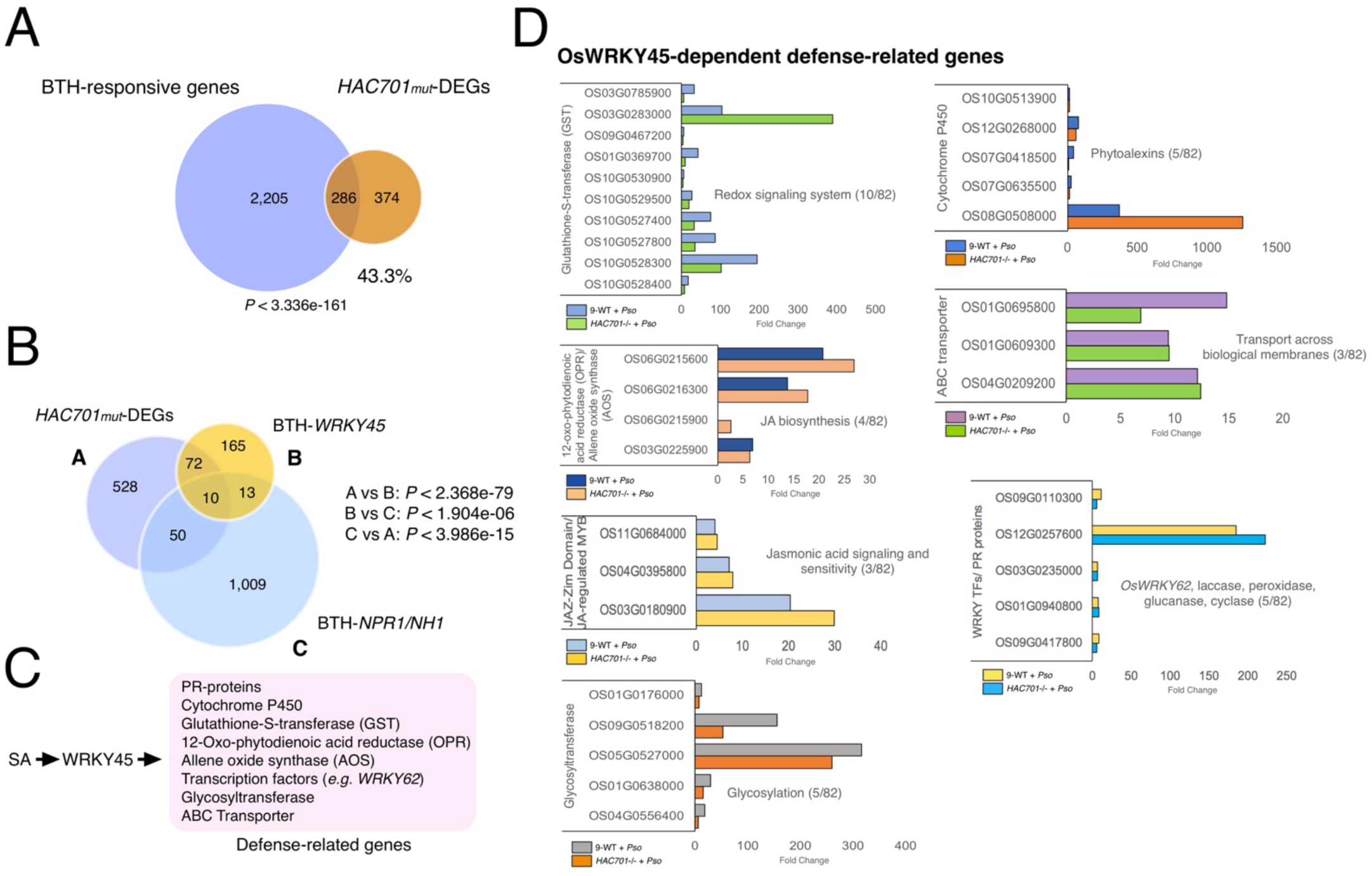
*hac701* phenocopies *OsWRKY45* overexpression transgenic plants. (*A*) Overlap analysis of differentially expressed genes (DEGs) in *hac701* background with benzothiadiazole (BTH)-responsive genes (*i.e.* BTH is a salicylic acid analog) in wild type rice (Table S6, S7). Significance value of the overlap data was tested using the hypergeometric distribution test. (B) Overlap analysis of DEGs in *hac701* background with BTH-inducible and *OsWRKY45*- dependent or *OsNPR1/NH1*-dependent genes (Table S8, S10). Significance values of the overlap data were tested using the hypergeometric distribution test. (*C*) *OsWRKY45*- dependent defense pathway components. (*D*) Gene expression of *OsWRKY45*-dependent defense genes in 9-WT and *hac701*.

Surprisingly, the upregulation of *WRKY45* expression in *hac701* did not result in any visible morphological defects (Fig. S1G). This indicates that *WRKY45* does not pose growth penalties, at least in this case where there is no pathogen applied.

### *WRKY45* promoter contains CRE motif

Rice HAC701 and two others, HAC703 and HAC704, belong to the CBP [<u>C</u>yclic adenosine monophosphate response element-binding protein (CREB) <u>B</u>inding <u>P</u>rotein] family group of proteins that is known to bind CREB transcription factor through its KIX domain (7, 37). Therefore, we examined *in silico* the members of rice CBP family for the presence of KIX domains using a KIX domain database (38). Our results showed that only HAC701 and HAC703 proteins contain the highly conserved KIX domain (Fig. S8). Although we have not identified CREB-like transcription factor candidates in rice so far, we further examined the DNA motifs in the 1kb promoter regions of the 660 *HAC701* mutant-DEGs in *Pso* that might potentially bind to KIX-CREB-like transcription factor. Our DNA motif analyses result showed 49 significantly enriched motifs in promoters (Table S13; E-value <0.05). We then used these motifs to compare against a database of known motifs and found a CRE motif when the query sequence CGRCGRCG (E-value<1.5e-27) was used (Fig. S9A). The CRE motif is a well-known sequence motif in animals usually bound by KIX-CREB transcription factor complex (39). We then searched for genes that contain the CRE motif and found that only five genes (0.76%) contain full CRE motif (Fig. S9B; Table S6). To our surprise, *WRKY45* promoter contains the full CRE motif that could possibly be targeted via KIX domain and CREB-like transcription factor-dependent regulation in rice system (Fig. S9C). These data show that *WRKY45* promoter could possibly be *cis*-regulated by HAC701 through its KIX domain, while the occurrence of CRE motif is not necessarily associated with other HAC701-regulated genes.

### Genome-wide investigation of histone modifications and enriched gene targets in *HAC701* **mutant**

The CREB-binding protein (CBP) is also known to regulate gene expression by acetylating histone tails in both animals and plants (40). Thus, we performed chromatin immunoprecipitation coupled to sequencing (ChIP-seq) analysis on histone markers associated with active and repressive chromatin to examine the landscape of chromatin modifications in *HAC701* mutant. We performed ChIP-seq analyses of H3K9 acetylation (H3K9ac), H3K27 acetylation (H3K27ac), H3K9 di-methylation (H3K9me2), and H3K9 tri-methylation (H3K9me3) on untreated 9-WT and *9-12b^-/-^* mutant (Fig. S10; Table S14). Metaplot profiles of these histone modifications were generated for genes, transposable elements (TEs), and simple repeats (Fig. S11; Fig. S12). The enrichment profiles show that both H3K9ac and H3K27ac accumulated in genes mainly along the transcriptional start site (TSS) and tapering through the gene body regions. In contrast, we found that H3K9me2 and H3K9me3 were highly depleted in the gene TSS region (Fig. S11A). As expected, H3K9 and H3K27 acetylation marks were depleted in TEs, while H3K9 di- and tri-methylation marks showed enrichment (Fig. S11B). In addition, enrichment analysis of simple repeats showed similar profiles with TEs in both acetylation and methylation histone marks, although simple repeats tend to be highly enriched in H3K9me2 at a narrow set of genomic loci (Fig. S12). These results showed that our ChIP-seq profiles of histone modifications were largely consistent with the previous report on rice ChIP-seq analysis (41).

The observed low enrichment of H3K9ac in *9-12b^-/-^* mutant is a probable indication of mutation on *HAC701* gene (Fig. S11A). Therefore, we further analyzed the significantly enriched or depleted regions in *9-12b^-/-^* by taking the enrichment ratio of 9-WT over *9-12b^-/-^* (wild type/*hac701*) in H3K9ac, H3K27ac, H3K9me2, and H3K9me3 modifications. These analyses yielded 265 peaks in the H3K9ac fraction (Fig. 4A; Table S14). From the metaplot analysis, it showed that *9-12b^-/-^* has depleted H3K9ac enrichment in the 265 regions identified containing putatively 263 genes (Fig. 4A; Table S15), which include *HAC704* and *HAC701* loci and a mediator gene, *MED11* (Fig. 4D; Fig. S13A). These 265 regions were not associated with changes in other modifications in *9-12b^-/-^* mutant when compared to 9-WT (Fig. 4B). These results suggest that HAC701 acetyltransferase may have a specificity to H3K9 site. It also suggests that HAC701 probably autoacetylates its own promoter including another member of CBP family, *HAC704*.

**Figure 4.**
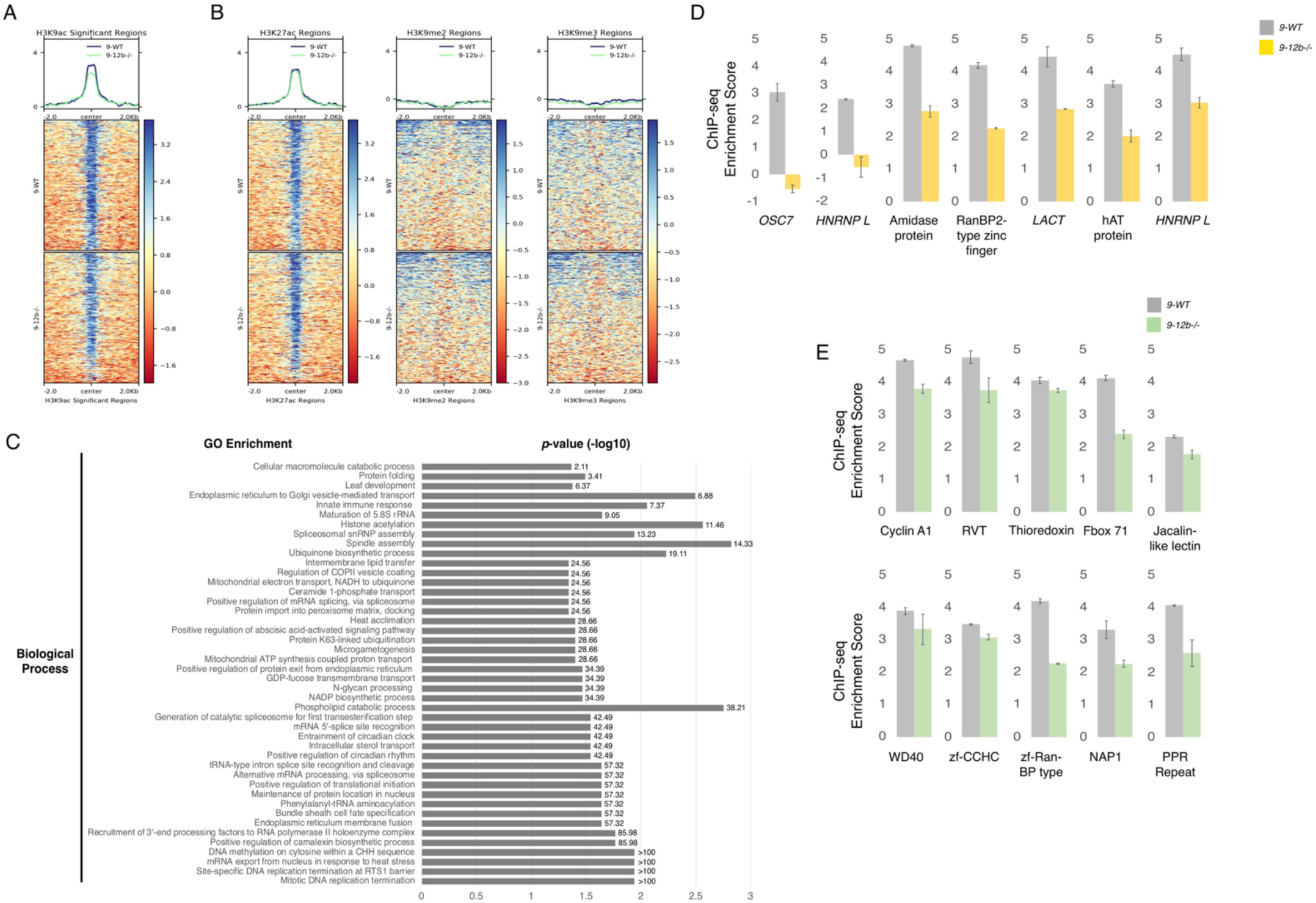
Genome-wide H3K9 acetylation is depleted in *hac701*. (*A*) Heatmap showing the accumulation of H3K9 acetylation on 265 peaks by taking the enrichment ratio of 9-WT and *9-12b^-/-^* data (Wild type/mutant) (Table S15). (*B*) Heatmaps showing the accumulation of H3K27 acetylation, H3K9 di-methylation, and H3K9 tri-methylation on 265 peaks by taking the same enrichment ratio as (*A*). Heatmaps in (*A*) and (*B*) are representatives of two biologically independent data showing similar enrichment results and number of peaks identified. (*C*) GO enrichment analysis showing only the “Biological Process” of the 263 genes found in the 265 peaks (Table S15). Numbers in enrichment indicate Fold Enrichment Value using Fisher’s Exact test with results for uncorrected *P*<0.05. (*D*) Enrichment scores of seven representative genes showing depletion of H3K9 acetylation in *hac701*, *9-12b^-/-^*. (*E*) Enrichment scores of 10 representative putative decoy/sensor genes (42) showing depletion of H3K9 acetylation in *hac701*, *9-12b^-/-^*. Scores in (*D*) and (*E*) are averages of two biologically independent ChIP-seq replicates with bars showing standard deviation (SD).

We then analyzed the GO enrichment features of the putative 263 genes found in these 265 regions with H3K9ac changes (Fig. 4C; Table S15). The results indicate that these genes function in processes involved in the flow of genetic information, DNA methylation in the CHH-context, circadian rhythm, organellar transport, cell cycle and reproduction, ubiquitination, and in a number of biosynthetic processes. It is also perhaps not surprising that histone acetylation was also enriched. In addition, enrichment of genes involving environmental responses included heat stress and innate immunity were observed, in which about 9% (23/263) were possibly involved in defense responses. A large fraction of these defense genes were identified as candidate sensors or decoys (e.g. Cyclin A1, Thioredoxin, Jacalin-like lectin, *etc.*) that may act as negative regulators of rice effector-triggered immunity (ETI-immunity) (42) (Fig. 4E; Fig. S13B). The distribution of these 265 regions in H3K9ac ChIP revealed that a large portion were found in genes (73.58%) followed by promoters (18.11%) (Fig. S14). It is also interesting to note that H3K9ac in these identified regions was found in transposable elements (7.55%). Overall, these results demonstrate that HAC701- dependent acetylation of H3K9 histone site might modulate transcription of decoy genes that are monitored by R proteins in rice immunity. Additionally, H3K9ac assessed in different regions of the rice genome showed that it is found mostly in genes and promoters with high similarity in the previous study (41).

We proceeded by analyzing the general distribution of H3K9ac, H3K27ac, H3K9me2, and H3K9me3 peaks in intergenic and intragenic regions of rice genome in 9-WT and *9-12b^-/-^* mutant (Fig. 5A). A large proportion of H3K9ac and H3K27ac were concentrated on exonic and intronic regions of the gene, while *HAC701* mutation resulted in the increase of enrichment of H3K27ac genome-wide on all regions of the rice genome (Fig. 5A). It is also worth mentioning that loss of *HAC701* increased the level of enrichment of these methylation modifications in intergenic regions. These data suggest that the loss of *HAC701* leads to an increase of a counter acetylation modification, H3K27ac, on a genome-wide scale.

**Figure 5.**
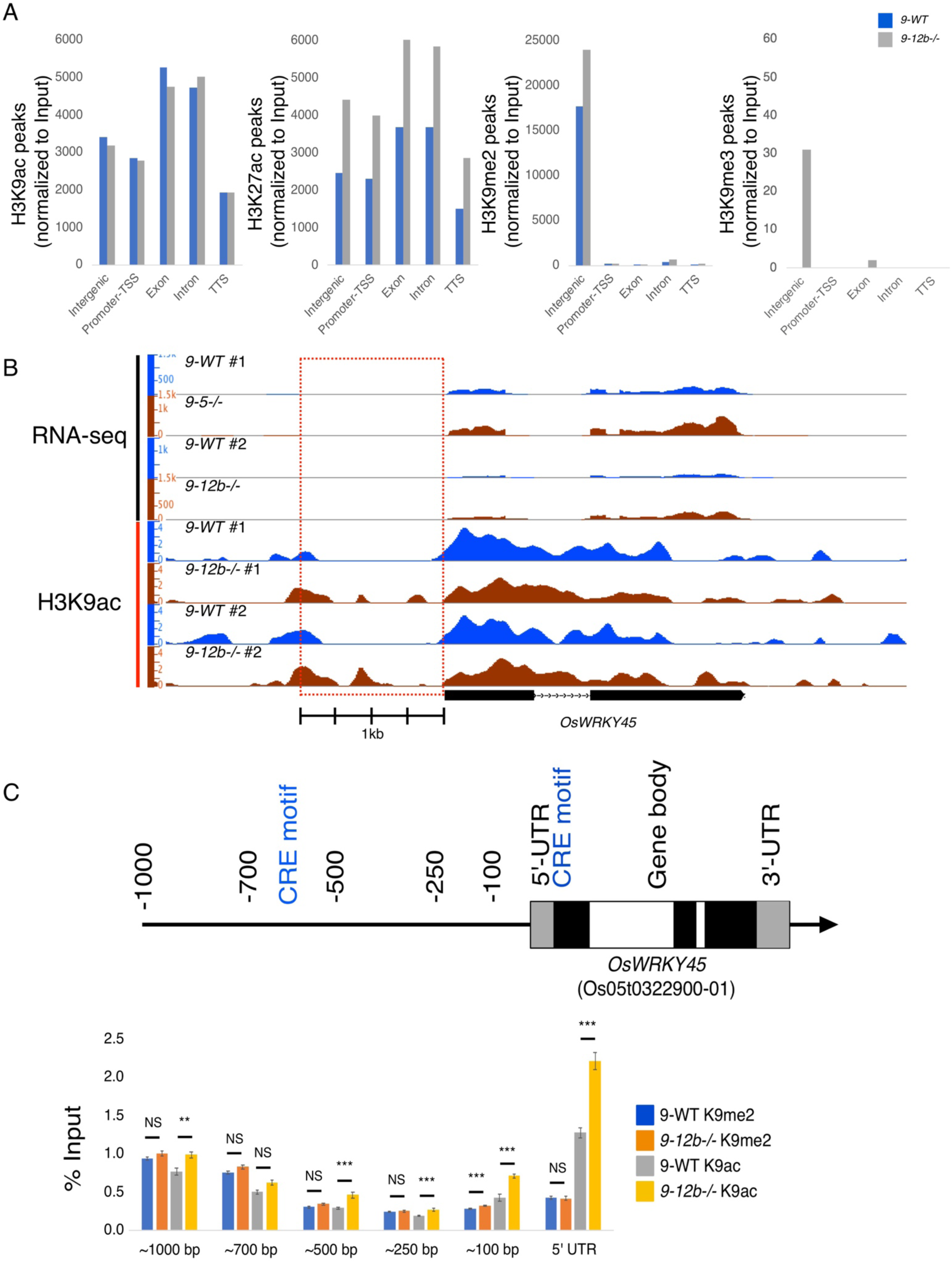
*OsWRKY45* loci is enriched in H3K9 acetylation in *hac701*. (*A*) Enriched peaks on histone modifications positioned to genomic locations (Intergenic, Promoter-TSS, Exon, Intron, and TTS) in 9-WT and *hac701*, *9-12b^-/-^*. All data were normalized to input. (*B*) *OsWRKY45* depth graph of RNA-seq and H3K9 acetylation ChIP-seq showing the number of reads in RPKM (Reads Per Kilobase of transcript, per Million mapped reads) in 9-WT and *hac701*, *9-12b^-/-^*. The red dotted box shows the 1kb upstream region of *OsWRKY45* with arrow indicating the direction of transcription. (*C*) Upper panel: Gene map of *OsWRKY45* with arrow showing the direction of transcription, gray bars are 5’ and 3’ untranslated regions (UTRs), black bars are exons, and white bars are introns. Lower panel: 9-WT and *hac701* (*9-12b^-/-^*) mutant ChIP-qPCR showing enrichments of H3K9 acetylation and H3K9 di-methylation in *OsWRKY45* locus. The significant difference in enrichment is computed using two-tailed Student’s *t*-test where asterisks: \*\*\**P<*0.01, \*\**P<*0.05, and NS means Not Significant.

To explain the disease resistance phenotype of *hac701* through WRK45-dependent immune pathway, we investigated the genomic locus of rice *WRKY45* (Fig. 5B). The upregulation of WRKY45 was accompanied by H3K9ac enrichment along the 1kb upstream region of *WRKY45* gene in *9-12b^-/-^* (Fig. 5B, C). As the CRE motif was specifically enriched in the promoter region of *WRKY45* (Fig. S9C), we compared the enrichment of H3K9ac and H3K27ac in *9-12b^-/-^ WRKY45* promoter. We found that H3K9ac, but not H3K27ac, was highly enriched in this regulatory region (Fig. S15), while H3K27ac of *WRKY45* gene body was less pronounced than H3K9ac. Overall, our ChIP-seq data are consistent with RNA-seq data in showing that mutation in *HAC701* gene potentiated *WRKY45* locus by regulating the enrichments of histone acetylation and methylation modifications on *WRKY45* promoter. We also found that the CRE motif of *WRKY45* is modulated by H3K9ac and perhaps other modifications, including those that are analyzed here.

### Systemic gene expression in rice-*Pso* pathosystem

To explore the nature of systemic gene expression in rice-*Pso* pathosystem, we performed RNA-sequencing on systemic or distal tissues of 9-WT and *hac701*. The tissue samples were collected together with infected samples (local samples) 72 hours after *Pso* infection without further bacterial treatment. Genome-wide transcriptome analysis in systemic 9-WT samples showed that 24 genes were differentially expressed, and among them were eight genes that were common to both local and systemic tissues (Fig. S16A; Table S16). Comparative analysis also showed that although there were substantial expression changes in systemic tissues, the number of DEGs was highly reduced in systemic tissues than in local tissues. Among the eight DEGs, five of them showed altered transcriptional expression from local to systemic tissues (Fig. S16B). These genes were mannose-specific jacalin-related lectin/*OsJAC1* (Os12g0247700) (43), putative cytochrome P450 (Os09g0275400) (44), ATPase (Os07g0187400), GDP-L-galactose phosphorylase (Os12g0190000) (45), and terpene synthase (Os04g0344100), some with putative functions in response to pathogen infection or stress. These results indicate that these genes were mostly activated locally in the infection site and that any altered expression in distal tissues may facilitate systemic form of resistance. To compare the effect of *HAC701* mutation on the number of DEGs, we analyzed the MA-plots of systemic tissues of 9-WT and *hac701*. Our results showed that the number of DEGs with significant expression (*i.e.* those that are in red) was diminished in the mutant background implicating a possible role of *HAC701* in systemic signaling in rice-*Pso* pathosystem (Fig. S17). Overall, these results indicate that systemic tissues in *Pso*-challenged 9-WT plants have augmented transcriptional gene expression that possibly aims to potentiate distal tissues on the onset of secondary pathogen attack. This also indicates that *HAC701* might potentially regulate systemic defenses through an unknown mechanism at distal non-infected site in preparation for future infection episodes.

## DISCUSSION

Histone acetyltransferases or HATs and their complexes are involved in various biological processes in the cell. Currently, studies have been focused on the roles of HATs specifically members of the CREB-binding protein (CBP) family in regulating substrate specificity and enzymatic activity such as acetylation (40). However, its involvement in molecular signaling pathway leading to modulation of plant immunity remains to be explored. Therefore, it is expected that the analysis of plant CBP will facilitate a broader understanding of regulation of gene expression and protein function in plant immunity pathways.

In this study, we generated CRISPR/Cas9 mutants of the rice *HAC701* gene, a member of the CBP family of HATs, since our results indicated its involvement in rice immunity. By inoculating a rice compatible bacterial pathogen, *Pseudomonas syringae* pv. *oryzae* (*Pso*), we showed that *hac701* are resistant to *Pso* proliferation and also showed resistance tendency when infected with *Magnaporthe* (rice blast). These indicates that *HAC701* negatively regulates rice immunity responses to *Pso* and *Magnaporthe* pathogens. When we analyzed our transcriptome data, it showed that this negative regulation of immunity by *HAC701* is attributable to suppression of a major immunity pathway in rice probably through WRKY45. In *hac701*, *WRKY45* gene expression is upregulated and it seemed to be in a potentiated state enabling a further increase of expression upon bacterial infection. In addition, both morphological phenotypes and target genes in *hac701* mimic those in *WRKY45-ox* transgenic lines possibly indicating a genetic interaction of *HAC701* and *WRKY45* genes. Lastly, we showed in our ChIP-seq analysis that the promoter of *WRKY45* in *hac701* is enriched in H3K9ac coinciding with a *cis* regulatory CRE motif that might be responsible for *WRKY45* upregulation and control.

In Arabidopsis, the p300/CBP acetyltransferase gene, *HAC1*, has been shown to positively regulate defense priming only in repetitively abiotic-stressed plants (46). *hac1-1* did not show any immunity related responses prior to abiotic stress applications indicating that Arabidopsis *HAC1* requires several abiotic stress stimulations to activate its immunity. However, this is not the case in *HAC701* in rice. We showed that the rice *HAC701* did not need abiotic stress stimulation to be transcribed and to effect pathogen and disease specific phenotypes (Fig. 1; Fig. S2; Fig. S3). Rather, our results are consistent with the disease phenotype exhibited by Arabidopsis mutant of GCN5, an acetyltransferase member of the SAGA transcriptional coactivator complex catalyzing the H3K14 modification site and at the same time influencing the acetylation of H3K9 (47). The *gcn5* showed upregulation of SA-mediated immunity resulting in resistance against *Pseudomonas syringae* pv. *tomato* infection. In our study, we found that WRKY45-dependent pathway confers resistance to *hac701*. Consistent with the view that the rice hormone defense network does not support a dichotomous role of salicylic acid (SA) and jasmonic acid/ethylene (JA/ET) phytohormones in regulating rice immunity (48), we found no evidence in this study that SA-mediated immunity played a central role in *hac701* resistance phenotype. However, we found JA-related differential gene expression as components of WRKY45-dependent defense-related genes (Fig. 3D). Indeed, JA biosynthesis is known as a downstream target of rice WRKY45 (49). In rice immunity model, both SA and JA/ET are considered effective in defense responses against (hemi) biotrophic and necrotrophic pathogens (48, 50, 51). While SA-mediated *AtNPR1* regulates majority, if not all, of Arabidopsis immune responses (52), *OsWRKY45* and *OsNPR1/NH1* are mostly independent of each other (34; Fig. S18). Although, our results suggest that *hac701* DEGs partially overlapped with NPR1/NH1-dependent pathways (Fig. 3B). Given that the rice system has a constitutive high levels of SA compared to most plants including Arabidopsis and tobacco, and pathogen applications did not increase its basal SA level (53, 54), our results are consistent with this report that SA in rice may not directly act on pathogens, rather, it may function to potentiate endogenous defense pathways in which *WRKY45* plays the central role. *OsWRKY45-ox* showed strong resistance against blast pathogen, even though lacking the constitutive expression of defense genes prior to blast infection (34). It could be that disruption of HAC701-dependent pathways through *HAC701* mutation led to activation of WRKY45-dependent pathways in an already SA-potentiated rice network, although this needs to be further examined in detail. Our results are also consistent with the disease phenotype presented by non-sense mutation in *Med15* gene, a subunit of Mediator complex (55). The *med15b.D* wheat mutants were resistant to stem rust, and the authors attribute a part of this to segmental coregulation in which a certain portion of the chromosome containing R genes (i.e. specifically NLR) was differentially expressed in wild type compared to mutant. While we did not observe any form of segmental coregulation in our analysis of *hac701*, we found that mutations in *HAC701* resulted in downregulation of ETI components notably the decoys/sensors (Fig. 4E).

*OsHAC701*, *AtHAC1*, *AtGCN5*, and *TaMed15*, play a central role in regulating the transcriptional machinery. They likely do this regulation by associating themselves and their complexes to RNA polymerase II and transcription factors as shown by their homologous counterparts in animals (4, 14, 56, 57). This is possible because they are multidomain genes that have the capacity to form and maintain large complexes. Os*HAC701*, *AtHAC1*, and *AtGCN5* are also involved in regulating abiotic stress responses suggesting that the complexity and specificity of their activities are dependent on their interacting factors that recruit them to the biological pathway itself (6, 7). Furthermore, majority of these genes, *OsHAC701*, *AtHAC1*, and *TaMed15*, encode the KIX domain, a docking site for transcription factors reported to be involved in plant immunity (58).

In line with the above results, we propose that HAC701 is a negative regulator of WRKY45-dependent defense pathway in rice (Fig. S18). We suggest here two possible mechanistic insights into HAC701 regulation of WRKY45 that need further experimental evidence. First, similar to Med15, the HAC701-mediated immunity may require a loss or truncation of HAC701 in regulating the transcription of *WRK45* locus. Given that the promoter of *WRKY45* contains a rare CRE motif (Fig. S9), we assume that an unidentified CREB-like transcription factor with regulatory potential is interacting with HAC701 through its KIX domain (Fig. S8) and with the CRE motif of *WRKY45* promoter. This interaction may also include other protein interactors specific to *WRKY45* locus regulation. From our results, mutation of *HAC701* is enough to potentiate *WRKY45* and further addition of *Pso* enables *hac701* to mount higher *WRKY45* expression compared to 9-WT. These suggest to us that *WRKY45* locus is poised for activation as suggested by our ChIP-seq analysis of untreated *hac701* (Fig. 5). However, the H3K9 acetylation we observed in *WRKY45* promoter might not be catalyzed by HAC701 itself in *hac701*, but may come from other CBP family members (Fig. 1A; Fig. S1H) or other HATs. Although, we do not have ChIP-seq data for pathogen inoculated *hac701*, our RNA-seq data suggest that the poised *WRKY45* through *HAC701* mutation is enough to activate downstream defense likely through *WRK45*-dependent signaling. Second, the decoys/sensors component that we found may act as negative regulators as predicted by the Decoy model (Fig. 4E) (42, 59). It could be that HAC701 maintains the acetylation of these decoys/sensors to equilibrium. Any disturbance in their expression could possibly lead to activation of R genes monitoring these decoys/sensors, which could in effect activate WRKY45-dependent immunity. Indeed, we found seven R genes upregulated in *HAC701* mutation treated with *Pso* (Table S17). It is of course possible that these two mechanisms are occurring simultaneously in conferring resistance, although finer details of these proposed mechanisms are still unknown.

As HAC701 is a global transcriptional regulator, we found that it regulates a myriad number of biological processes underscoring its importance in genome and epigenome control (Fig. 4C). Among the enriched GOs, we found histone acetylation and innate immune responses were affected in *hac701*. These suggest to us the intricate network in HAC701- dependent regulation of immunity through its scaffolding and catalytic activity in the cell. It is possible that any misregulation in *hac701* could lead to genome-wide functional compensation events by activating other proteins acting on the same pathway as HAC701. This compensation functionality has been shown in a PHD-finger protein, Enhanced Downy Mildew 2 (EDM2), in NLR (i.e. a type of R protein) protein expression control as it affects plant fitness (60). Therefore, it is also likely that the expression regulation of *WRKY45* locus as well as the decoys/sensors may be subjected to compensatory regulation in the case of *HAC701* mutation.

The heterochromatic marker, H3K9me3, is common in animals, but rarely found in Arabidopsis and in plants in general (61). Indeed, our analysis did not find the peaks in 9-WT ChIP-seq, although we found 33 peaks in the *HAC701* mutant (Table S14). Most H3K9me3 enrichments were found in intergenic regions and in only two genes with intragenic enrichments. Table S18 also shows genes nearest the H3K9me3 enrichment in the intergenic regions. These results suggest that H3K9me3 is maybe a rare modification and is possibly modulated by HAC701 presence and activity in certain genomic regions.

Trade-off between growth and defense in plants is a common occurrence and often considered inevitable (62, 63). However, we were surprised to find that *hac701* in the T2 up until T4 generations do not have readily visible penalties in height, tiller number, number of effective grains, and grain weight (Fig. S1D-I). Previous studies showed that *WRKY45-ox* had no obvious autoimmunity phenotype, although there was a slight decrease in height observed as compared to wild type in response to growth conditions (33, 34). In this study, WRKY45- dependent resistance against *Pso* was observed in *hac701* (Fig. 1C, Fig. S3). Furthermore, these *HAC701* mutants also showed resistant tendencies against *Magnaporthe* infection (Fig. 1D). *WRKY45-ox* treated with *Magnaporthe* showed resistance, while a BTH-induction in knockdown *WRKY45* mutants did not rescue the *Magnaporthe* resistance and remained susceptible (34). Together these suggest that morphological and disease phenotypes of *hac701* resembled that of *WRKY45-ox* indicating that *HAC701* mutation indeed upregulates *WRKY45* gene that prompts WRKY45-dependent defense pathway into action leading to resistance against bacterial and fungal pathogens. It could be that the potentiated state of *WRKY45* in *hac701* has modulated the expression of resistance from affecting these agronomic traits. Nonetheless, the mechanistic basis for this trade-off, if it truly exists, is not well understood and needs further investigation.

We would like to emphasize that at the moment we do not have strong evidence that HAC701 specifically catalyzes H3K9 acetylation. Although, the H3K9ac antibody used in this study detected H3K9ac decrease of numerous loci in *hac701* compared to 9-WT that are not present in H3K27ac, histone target-specificity of HAC701 still needs to be verified. As H3K9ac and H3K14ac were seemed co-regulated and were directly correlated to each other in regulating their common target genes (47), we assume that the H3K14ac in our *hac701* would show the same correlation as well.

HAC701 appears to be an ancient protein in the CBP histone acetyltransferase family clade (Fig. 1A). The fact that *HAC701* mutation is not lethal, suggests that other CBP members and arguably members of other histone acetyltransferase families might have redundant functions at some regulatory level. However, our results showing that *hac701* has enhanced disease resistance phenotype and reduction in acetylation demonstrate certain functional specificities through HAC701-dependent pathways. Also, the fact that *hac701* showed reduction in expressed genes at the systemic level (Fig. S17), indicates possible broader roles of HAC701 other than localized modulation of rice innate immunity.

## MATERIALS AND METHODS

### Biological samples and plant growth conditions

*Oryza sativa* ssp. *japonica* cv. Nipponbare (wild type) plants and mutant lines were grown in commercial soil (Kumiai, JA Okinawa) at 30°C day/25°C night temperatures under a 12-h-light/12-h-dark photoperiod in an incubator (BiOTRON, NK System). The lighting was supplied by white light at an intensity of 31,000 lx. Relative humidity was at 70%. For embryonic rescue, rice embryos were extracted from wild-type immature seeds 10-14 days after flowering. Embryos were grown into plantlets in Murashige and Skoog (MS) agar medium before being transferred to soil for further growth.

### Isolation and screening of *HAC701* mutant lines

To generate *HAC701* CRISPR/Cas9 knockout mutant lines, two sgRNA *HAC701*-specific target sites were obtained from CRISPR-P website and were used to synthesize primers for pRGEB31 (stable system) (64) (Table S1). Briefly, the vectors were digested with BSA I, while primers were phosphorylated and annealed to produce a DNA oligo duplex. The digested vectors were ligated to DNA oligo duplex using T4 ligase. CRISPR/Cas9 vectors were introduced into *Agrobacterium* EHA105 and rice calli were transformed using the standard *Agrobacterium*-mediated transformation procedure. For CRISPR/Cas9 T0 and T1 screening, leaf samples were collected and genomic DNA was extracted using a standard CTAB protocol. Then, PCR-RFLP assay utilizing BseLI restriction enzyme (Thermo Fisher Scientific) was used to detect the CRISPR/Cas9-engineered mutations on *HAC701* targets (Table S1). Positive lines detected by PCR-RFLP assay were further analyzed by sequencing for INDEL mutations using S2 and S5 forward primers (Table S1). Four DNA amplicons per line were cloned and sequenced to determine the zygosity of the lines. Biallelic mutations were found in a few T0 and T1 lines. First exon (Target 1) was initially targeted in *HAC701* gene using a CRISPR/Cas9 vector construct containing a single-guided RNA (S2) (Fig. S1A; Table S1). The isolation of the first generation (T0) *CRISPR Cas9-HAC701-S2* lines yielded 90 positive independent lines, and among these, randomly chosen representative lines were further genotyped to characterize the identified DNA mutation. PCR and RFLP assays resulted in the isolation of monoallelic lines characterized mostly by deletions and a few insertions (INDELS) on or surrounding the targeted site of sgRNA (S2) (Fig. S1B-C; Table S1). The T1 *HAC701-S2* generation as observed from seeds showed conservation of mutation directly from parental lines (Fig. S1B). Then, mutant lines were isolated using the same technology targeting the fifth exon (Target 2) of the *HAC701* gene using a construct that also contains a single guide RNA (S5) (Fig. S1D; Table S1). The *CRISPR-Cas9-HAC701-S5* (T0) lines generated 11 positive independent lines of which three were genotyped for verification of mutations (Fig. S1E). Second generation (T1) lines also showed conservation of mutations (Fig. S1E). Similar to *HAC701-S2* lines, genotyping showed INDELS in proximity to or on the target site.

### Phenotyping

Positive lines containing biallelic homozygous mutations were phenotyped for effective grain production by examining and counting the mature grains produced in each panicle. Tiller number was counted one month after flowering. Chlorosis phenotype was used as a proxy for any defect in photosynthesis and or apparatus. 1000-grain weight was measured on seeds oven dried for 30-days at 37°C. Comparison of height and general morphological structures were documented using photography.

### flg22 leaf disc assay

flg22 peptide, a well-known inducer of plant innate immunity, specifically of PTI was used to test which rice HATs respond to flg22 treatment. Leaf disc assay (modified from 65, 66) was performed on fully expanded 30-d-old wild-type leaf samples and treated with synthetic flg22 peptide (30-51 aa, Flic, *Pseudomonas aeruginosa*) (ADI, Inc.) at different time periods and concentrations. Briefly, leaves were cut into about 5 mm sizes and floated on the water for 24-h in growth chamber to remove the symptoms of wounding stress. Leaves were then treated with PAMP solution in water at 15 ml falcon tubes with rotation (Corning Science). After treatment, paper towel-dried leaves were frozen in liquid nitrogen.

### Pathogenesis assay

Seeds were surface sterilized and imbibed in sterile water in the dark for 72 h before sowing on MS basal medium (Sigma Life Science). After 10 days, the plantlets were transferred to soil and were grown for another 18 days until infection. *Pseudomonas syringae* pv. *oryzae* (*Pso*) (MAFF No. 301530, NIAS Genebank) was grown in Luria-Bertani (LB) broth (Sigma-Aldrich) at 28°C until OD = 0.2 and was resuspended in 10 mM MgCl_2_. The fourth leaf counting from the first true leaf was infected with *Pso* using needleless syringe injected from the lower surface of the leaf. *Pso* leaf infiltration was performed 10 cm from the tip of the leaf and was done 3x with approximately 1 cm space between the infiltration sites. Infected plants were temporarily maintained outside the growth chamber for 2 h to allow drying of the infected sites before returning to the chamber. Infected leaf samples were collected 3 days after *Pso* inoculation utilizing only the tissues comprising the spaces between the infiltration sites. These tissues were grounded in sterile 10 mM MgCl_2_ and the grounded tissue suspensions were transferred to LB agar medium for incubation. The infected local tissues were assayed with minor modifications as detailed in Liu*, et al.* (67). For log cfu per leaf disc measurement, the grounded tissue suspensions were serially diluted six times to be able to count with accuracy the *Pso* colonies on Luria-Bertani (LB) agar medium. *Pso* colonies were counted from 4^th^ until 6^th^ serial dilution in several independent and genotypically dissimilar *hac701* lines*, 9-5^-/^*, *9-12a^-/-^*, and *9-12b^-/-^*. The remaining samples of the fourth leaf were used for RNA-sequencing analysis. For blast assay, the fifth leaf of 25-day-old plants were used for infection of *Magnaporthe oryzae* (MAFF No. 101511, NIAS Genebank). Blast spores were incubated 10 cm from the tip of the leaf (1^st^ site) and at another site of the same leaf 10 cm from the first site (2^nd^ site). Infected plants were incubated for 10 days inside an incubator (BiOTRON, NK System) and the length of blast infection was measured.

### Gene expression analysis

Total RNA extraction was performed using RNeasy Plant Mini Kit (Qiagen) or Maxwell^®^ 16 LEV Plant RNA Kit (Promega). cDNA was synthesized using Primescript II 1^st^ Strand cDNA Synthesis Kit (Takara) according to manufacturer’s instructions. RT-qPCR assays were performed on three biologically independent samples or as indicated. RT-qPCR was performed using SYBR Premix Ex Taq II (Tli RNaseH Plus) (Takara) and was calculated following Pfaffl (68) by averaging the values relative to *ACT1* control gene. Primer sequences are listed in Table S19.

### RNA-sequencing

Total RNA was isolated from mock- and *Pseudomonas syringae* pv. *oryzae* (*Pso*)-treated 9-WT wild type and *hac701* with Maxwell 16 LEV Plant RNA Kit (Promega) run on the Maxwell 16 Instrument (Promega) and/or mirVana miRNA Isolation Kit (Invitrogen by Thermo Fisher Scientific). The *hac701* lines, *9-5^-/-^* and *9-12b^-/-^*, were lumped together as two independent biological samples during analysis. To remove the contaminating genomic DNA from RNA samples isolated using the mirVana miRNA Isolation Kit, RNA was treated with DNase I (RNA free) (Nippon Gene) following the manufacturer’s instructions. Samples were submitted to OIST Sequencing Center for RNA quality checking, library preparation, and paired-end mRNA-sequencing (PE mRNA-seq).

### Chromatin immunoprecipitation-sequencing

ChIP analysis of histone modifications in wild type and mutants were performed as follows: One-month-old mature leaves of wild type (9-WT) and *hac701* (*9-12b^-/-^*) were fixed in a fixation buffer (10mM Tris-HCl (pH 7.5), 50 mM NaCl, 0.1 M sucrose, 1 % formaldehyde) for 10 min, followed by quenching with 125 mM Glycine for 5min. Nuclei isolation was performed as previously described (69). Immuno-precipitation was performed for two replicates for each genotype (about 1 g tissue/IP) by SimpleChIP Plus Kit (Cell Signaling Technology) according to the manufacturer’s instructions. Anti-Histone H3 antibody (Abcam ab1791), Anti-acetyl Histone H3K9 antibody (Merck ABE18), Anti-Dimethyl Histone H3K9 antibody (Merck 05-1249), Anti-Trimethyl Histone H3K9 antibody (Merck 17-10242), Anti-acetyl Histone H3K27 antibody (Abcam ab4729) were used for IPs. Dynabeads M-280 Sheep Anti-Rabbit IgG (Invitrogen 11203D) or Anti-Mouse IgG (Invitrogen 11201D) was used for purification of chromatin-antibody complex. Precipitated DNA samples were sequenced by Hiseq 4000 in 150 bp paired-end mode in OIST Sequencing Center. For ChIP real-time PCR, we used the “Percent Input Method” (Thermo Fisher Scientific) with 33% input value to normalize the data for enrichment calculation. Three independent biological replicates were performed for each sample.

### Data analysis

For RNA-sequencing analysis, high quality reads were trimmed in order to remove sequencing bias and adapter effects. Trimmed reads were then mapped to the *Oryza sativa* spp. *japonica* genome (Os-Nipponbare-Reference-IRGSP-1.0) using Tophat (70, 71). Custom R scripts were used to generate the RNA count table necessary to analyze the differentially expressed genes (DEGs). Differential expression analysis was performed using DESeq2 package (72). DEGs were selected at Benjamini-Hochberg adjusted *p* value of < 0.01. To perform Gene Ontology (GO) analysis, gene lists were submitted to Enrichment Analysis (73) web tool of the Gene Ontology Consortium (74–76). To create the Venn diagram, gene sets were submitted to InteractiVenn web tool using *unions by list* to obtain overlapping components of input datasets (77). For ChIP-sequencing analysis, raw reads were trimmed and were mapped to *Oryza sativa* spp. *japonica* genome (Os-Nipponbare-Reference-IRGSP-1.0) using Bowtie (78). Read alignments were visualized using Integrated Genome Browser (IGB) (79). ChIP-enriched peaks were called by deriving a quantitative reproducibility score called “irreproducibility discovery rate” (IDR) (73). Comparative analyses and quantitation of enrichments between 9-WT and *hac701* (*9-12b^-/-^*) were analyzed using a suite of tools and utilities found in deepTools (80) and BEDTools (81). Peaks were positioned to genomic locations (intergenic, genes, transposable elements, simple repeats) and within genes (promoter, exon, intron, TSS, TTS). Table S14 showed the number of peaks identified for each histone modifications in 9-WT and *hac701* (*9-12b^-/-^*). Peak-to-gene assignment was performed using Homer (v4.11) (82). To create the phylogenetic tree of Arabidopsis and rice histone acetyltransferases, we used the UniProt amino acid sequences and reconstructed the phylogenetic relationships using Phylogeny.fr (83, 84). For KIX domain analysis, we used KIXBASE to perform multiple sequence alignment (38). To identify *cis*-elements on DNA sequences, we used cister prediction software that utilizes hidden Markov model (85). For motif analyses, we utilized the 1kb upstream promoter sequences of the sample genes. We used Discriminative Regular Expression Motif Elicitation (DREME) (86) to discover the significantly enriched motifs at a threshold of E-value<0.05 using shuffled input sequences as control. These enriched motifs were used as query sequences in Tomtom (MEME suite 5.2.0) to find similar motifs in published libraries. Libraries used were Eukaryotic DNA and Vertebrates (*in vivo* and *in silico*).

### Data visualization

Visualization of majority of the data was performed using Microsoft Excel for Mac 2011 (version 14.7.1) and R packages: heatmaply, functions from *gplots*, and *ggplot2*.

### Data Repository

For RNA-sequencing and ChIP-sequencing, raw data have been deposited in the DDBJ Sequence Read Archive under accession ID (pending upon submission of data). All other data can be found in the Supplemental Tables 1-19 in this manuscript.

## TABLE LEGENDS

None

## SUPPLEMENTARY FIGURE LEGENDS

**Figure S1.**
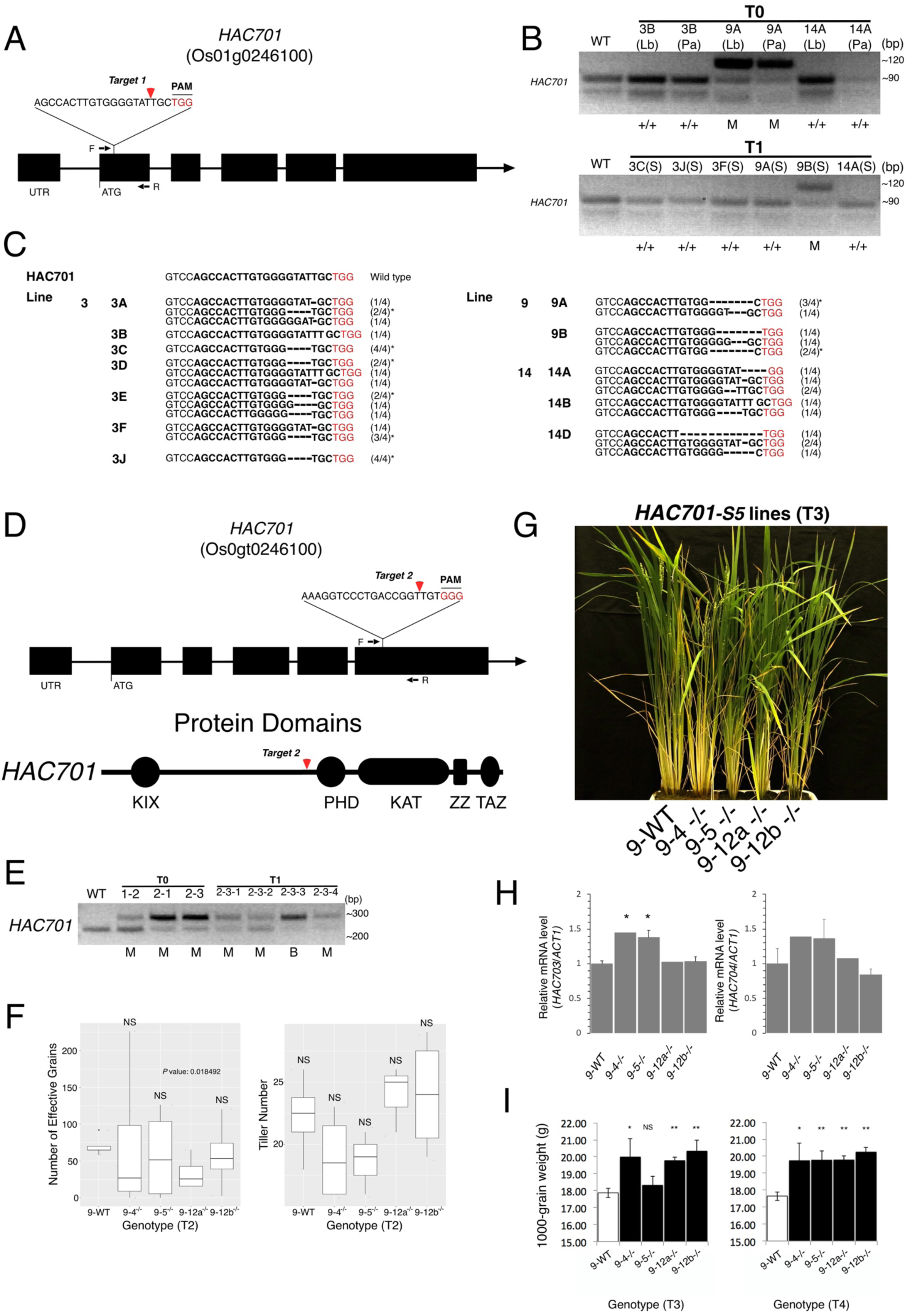
Characterization of mutations generated by CRISPR/Cas9 editing on the first and fifth exon of rice acetyltransferase gene, *HAC701*. (*A*) Schematic showing an sgRNA targeted to the first exon of *HAC701* gene. (*B*) PCR and RFLP assays of representative T0 and T1 generation lines from leaf blade (Lb), panicle (Pa), and seed (S) DNA samples. +/+ and M represent the zygosity of the line, where +/+ refers to wild type and M refers to monoallelic. (C) Alleles from 12 T0 generation lines identified by cloning and sequencing the PCR products from *HAC701* target regions using the primers found in Table S1. Similar line number indicates that lines came from the same callus. For each line, four DNA amplicons were cloned and sequenced and the fraction indicates the number of times the type of mutations were found in each line. In case not indicated, it means wild type. The asterisk (*) indicates the most common mutation found within and across different lines. (*D*) Upper panel: Schematic showing an sgRNA targeted to the fifth exon of *HAC701*. Lower panel: Schematic showing the protein domains of HAC701. (*E*) PCR and RFLP assays of representative T0 and T1 generation lines from leaf blade DNA samples. M, B represent the zygosity of the line, where M refers to monoallelic, and B refers to biallelic. T1 lines came from parental 2-3 line. (*F*) Phenotype of T2 lines showing effective grains and tiller number. (*G*) Images of T3 *hac701* lines compared to segregated wild type, 9-WT. (*H*) RT-qPCR levels of *HAC703* and *HAC704* genes in *HAC701* mutation line background. Samples were treated hydroponically with HDAC inhibitors (1µM Trichostatin A (TSA) and 100 µM Nicotinamide at final concentration) for three days. Bars represent standard error (SE); n= 3. The significant difference in transcription is computed using two-tailed Student’s *t*-test where asterisks: \**P<*0.01. (*I*) Phenotype of T3 and T4 lines showing grain weight.

**Figure S2.**
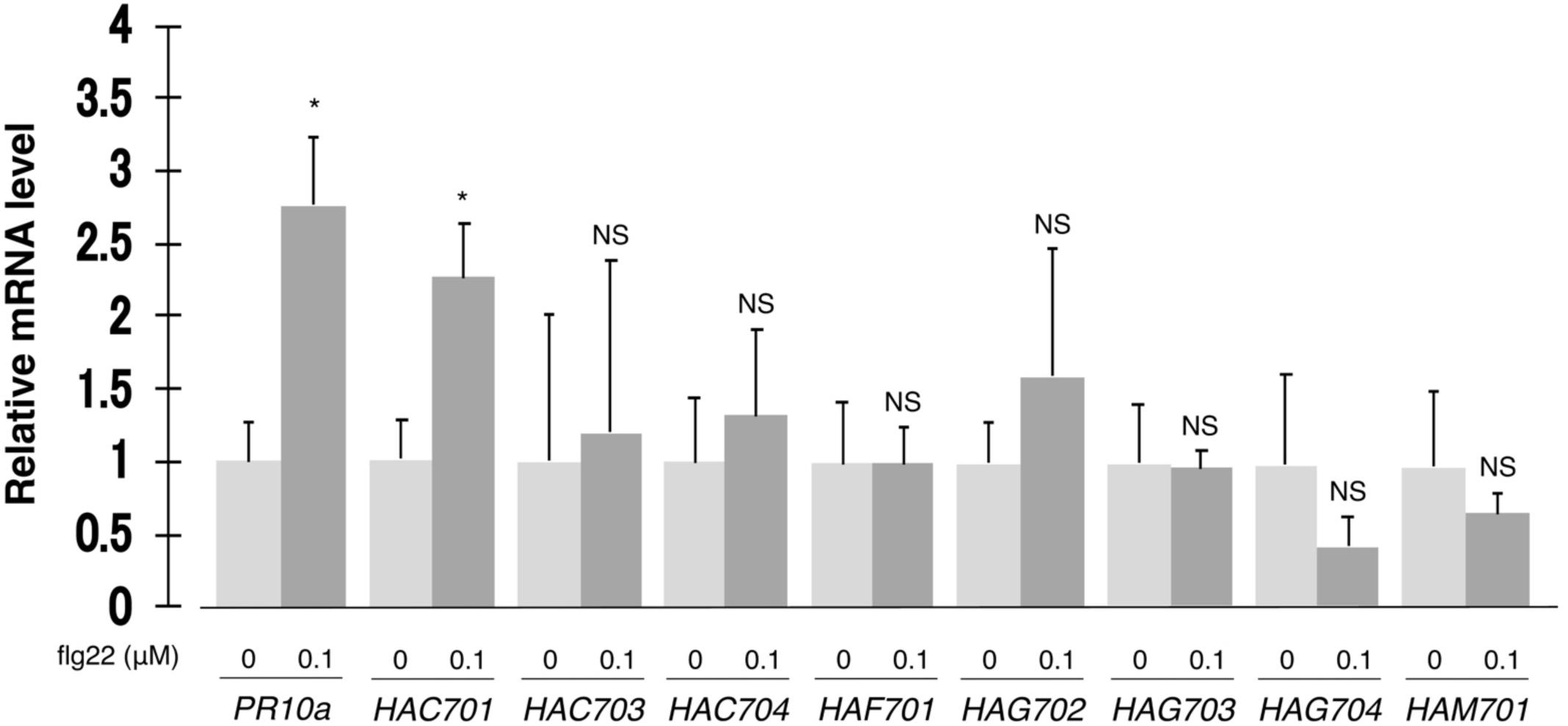
flg22 induced the expression of rice pathogenesis-related and histone acetyltransferase (HAT) gene, *HAC701*. Transcriptional levels of pathogenesis-related gene, *PR10a*, and of eight HAT genes upon flg22 induction at concentrations in µM units. Data shown are means ± SE; n= 3. The significant difference in transcription is computed using two-tailed Student’s *t*-test where asterisks: \*\*\**P≤*0.01, \*\**P≤*0.03, * *P≤*0.05.

**Figure S3.**
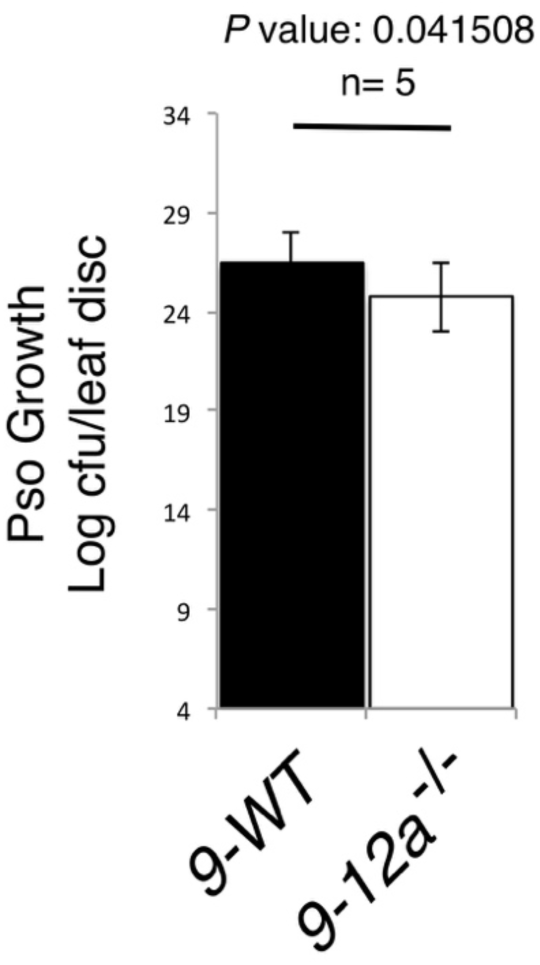
P*s*eudomonas *syringae* pv. *oryzae* (*Pso*) infection assay in T2 *hac701* line, *9-12a^-/-^*. Values from 4^th^-6^th^ serial dilutions were used for quantification. Segregated wild type line was used as a control. Bars represent the 95% confidence interval (CI) and compared to wild type using one-tailed Student’s *t*-test at *P*<0.05.

**Figure S4.**
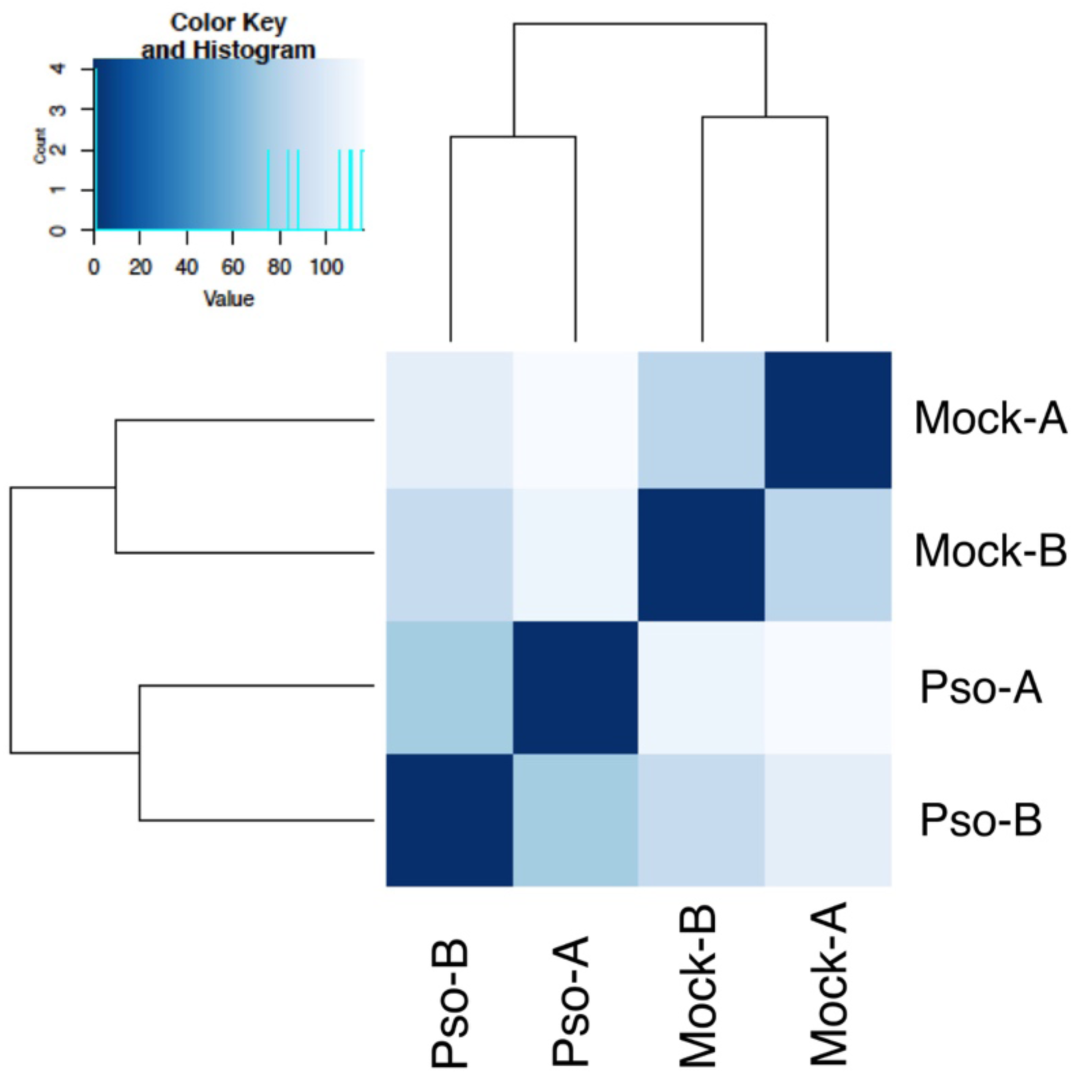
Sample distances of RNA-seq data after regularized-logarithm transformation (rlog). Heatmap showing the Euclidean sample distance matrix of mock- and *Pseudomonas syringae* pv. *oryzae* (*Pso*)-treated samples in two biologically independent replicates.

**Figure S5.**
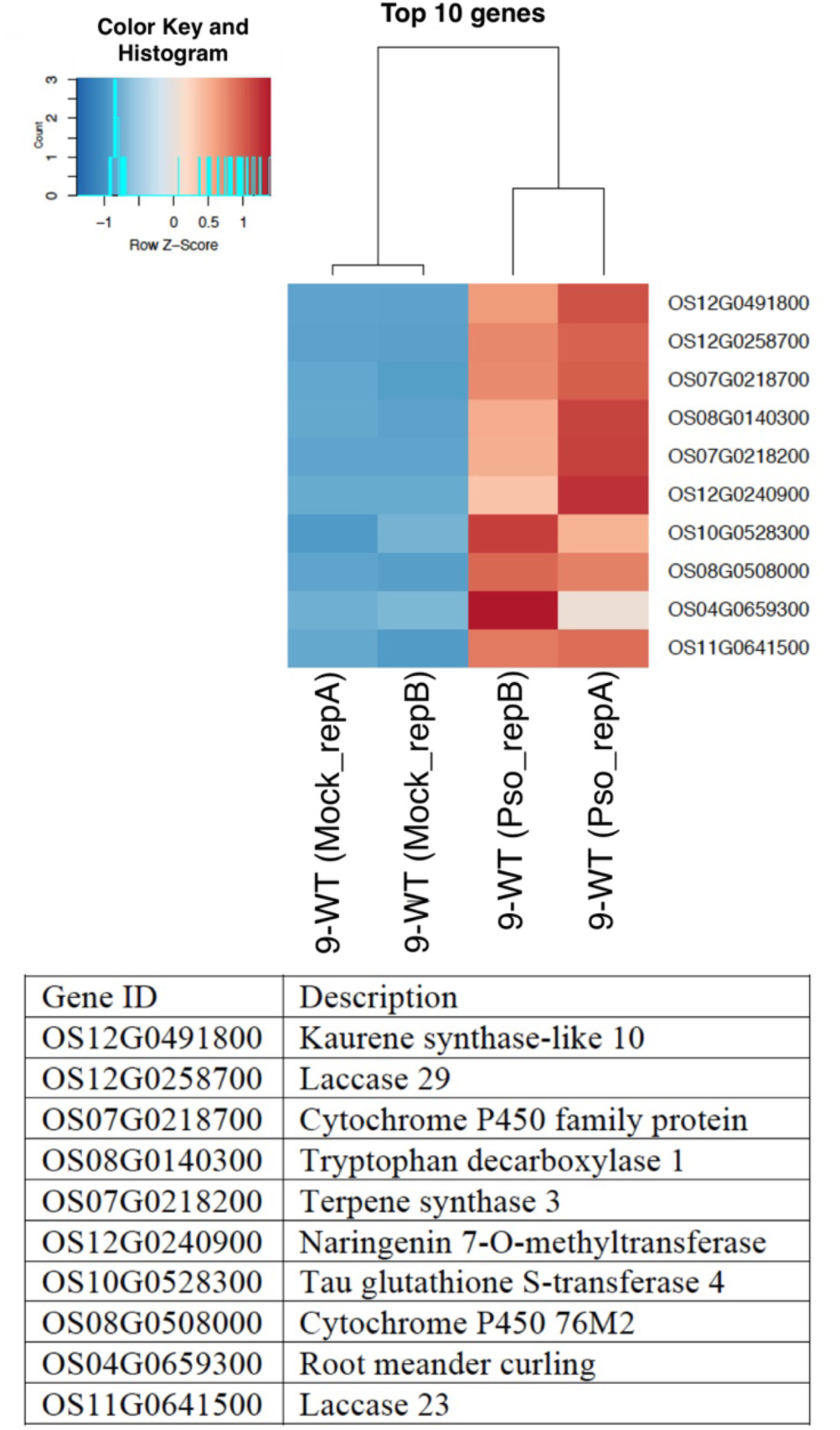
Top 10 genes that are most highly variable in mock- and *Pso*-treated RNA-seq samples. Gene descriptions were derived from Oryzabase: Integrated Rice Science Database (NBRP) and Rice Genome Annotation Project (NSF). Two biologically independent RNA-seq replicates are presented.

**Figure S6.**
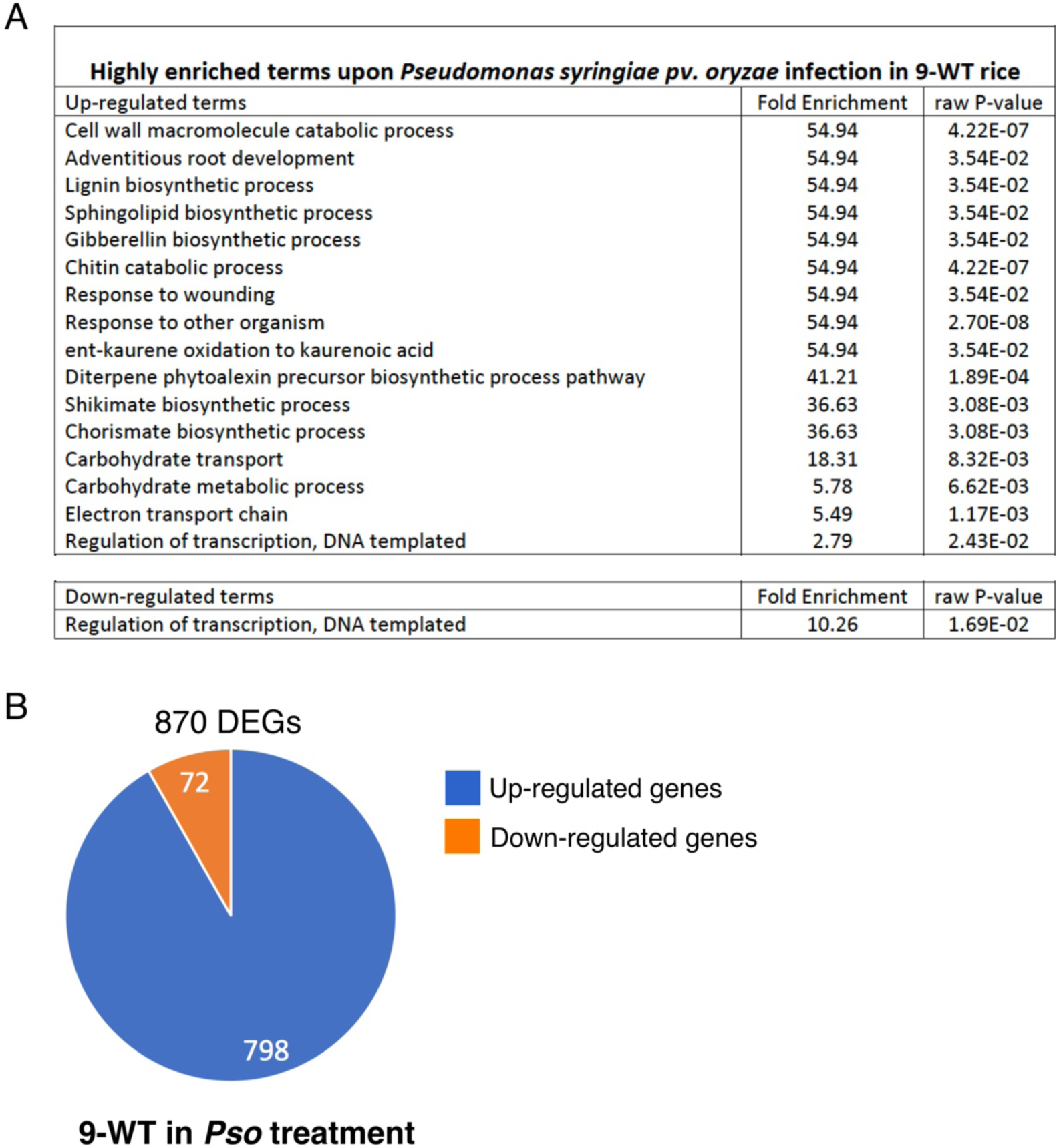
Rice-*Pseudomonas syringae* pv. *oryzae* (*Pso*) pathosystem. (*A*) Enriched pathways in the rice-*Pso* pathosystem. (*B*) Pie chart of the number of up- and down-regulated genes in 9-WT control 72h post *Pso* infection (Table S3). Two biologically independent RNA-seq data in mock and *Pso* treatments are presented where differentially expressed genes (DEGs) were selected at *P* adjusted value < 0.01.

**Figure S7.**
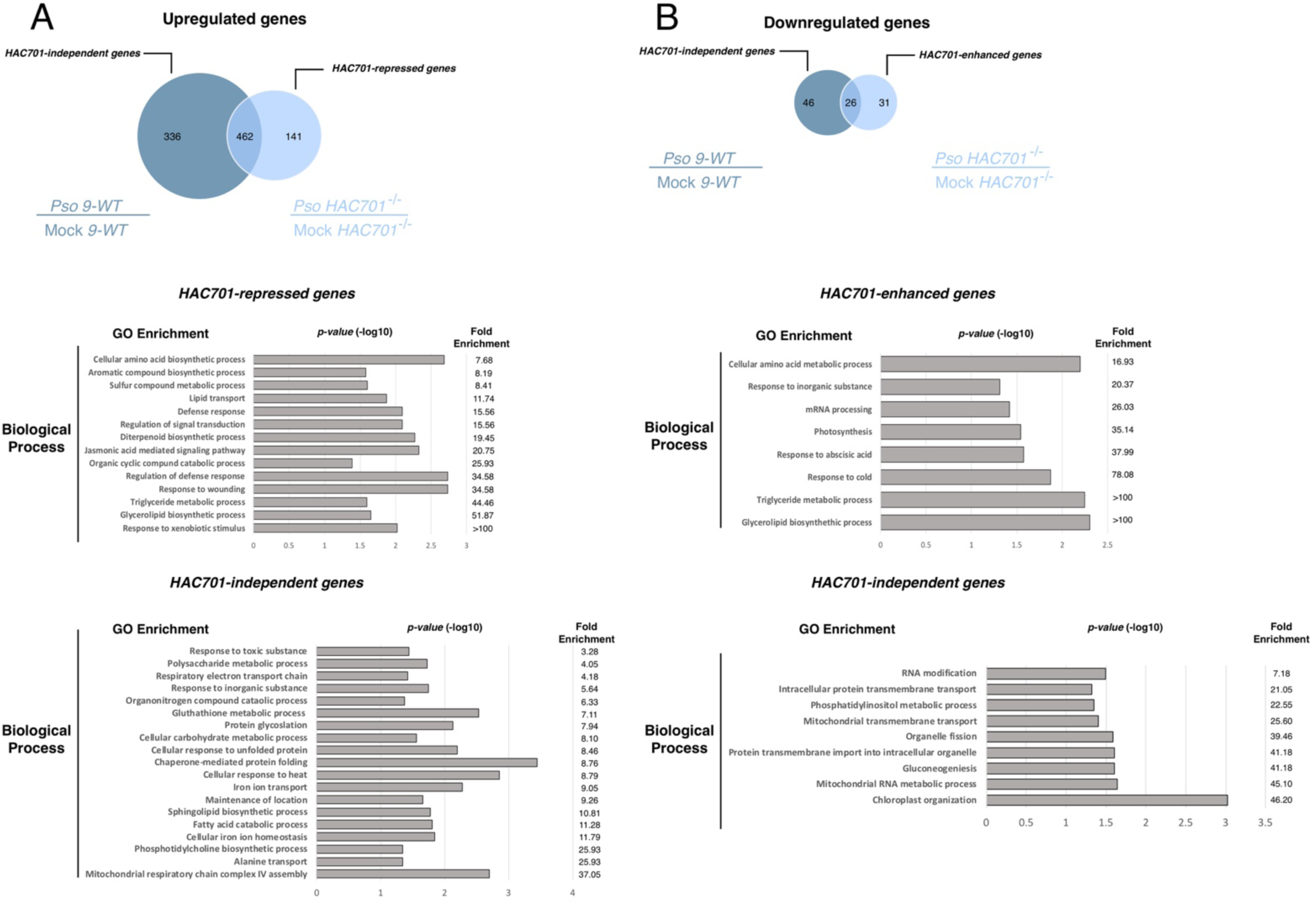
Features of differentially expressed genes (DEGs) in *Pseudomonas syringae* pv. *oryzae* (*Pso*)-treated 9-WT and *hac701* transcriptomes. Overlap and GO-analysis of upregulated (*A*) and downregulated (*B*) genes in *hac701* under *Pso* treatment (Table S3, S4). Transcriptome data were normalized using mock data sets for each genotype. GO enrichment analysis showing only the “Biological Process” of upregulated *HAC701*-repressed genes (141) and HAC701-independent genes (336), downregulated *HAC701*-enhanced genes (31) and HAC701-independent genes (46). Fold Enrichment Value uses Fisher’s Exact test with results for uncorrected *P*<0.05.

**Figure S8.**
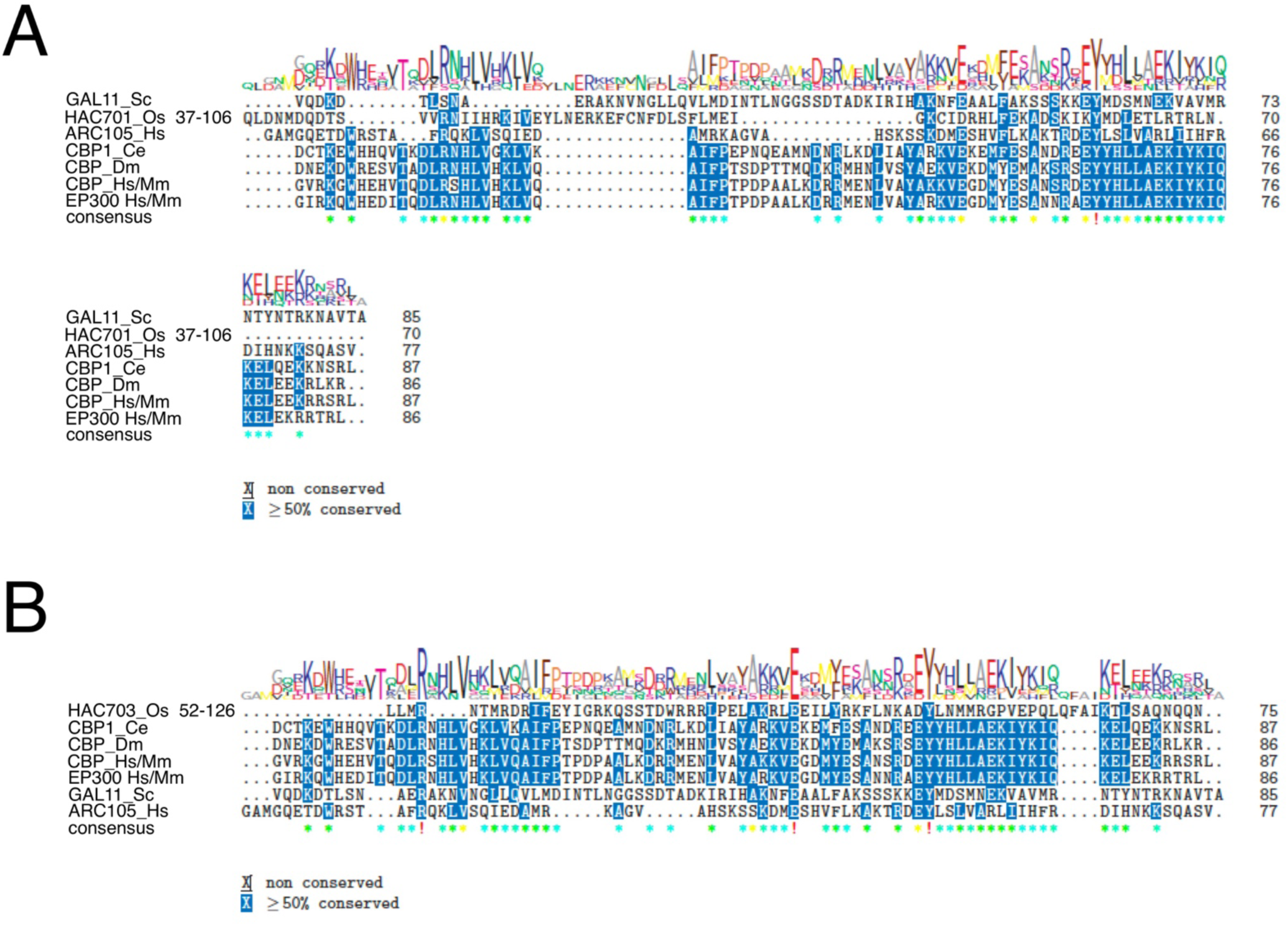
Alignment of KIX domain found in different organisms. KIX domain in HAC701 (*A*) and HAC703 (*B*) proteins. Sc: *Saccharomyces cerevisiae*, Os: *Oryza sativa*, Hs: *Homo sapiens*, Ce: *Caenorhabditis elegans*, Dm: *Drosophila melanogaster*, Mm: *Mus musculus*.

**Figure S9.**
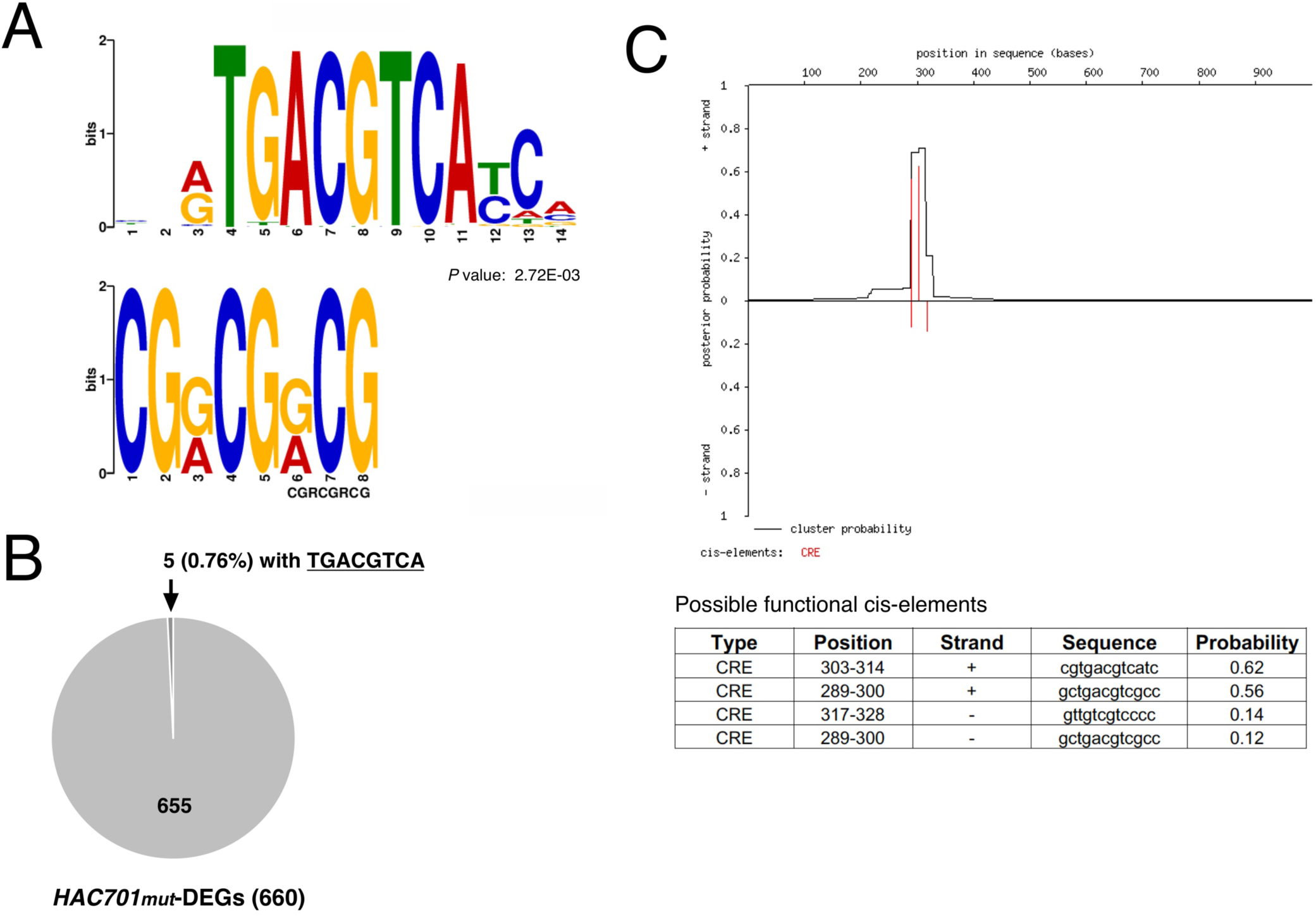
O*s*WRKY45 promoter contains a rare CRE motif. (*A*) A CRE motif (upper panel) matches significantly to the query motif CGRCGRCG (lower panel). The query motif was derived from discovered motifs (Table S13) in the 1 kb promoter regions of differentially expressed genes (DEGs) in *hac701* treated with *Pso*. (*B*) 5 genes out of 660 differentially expressed genes (DEGs) (0.76%) (Table S6) in the *hac701* contain the full CRE motif. Cister parameter settings: default settings were used. *(C)* CRE motif cluster found in 1kb upstream sequences of *OsWRKY45* start site. The red line indicates the probability of transcription factors binding to these CRE motif sites. CRE motif is found most probably on the direct strand (+).

**Figure S10.**
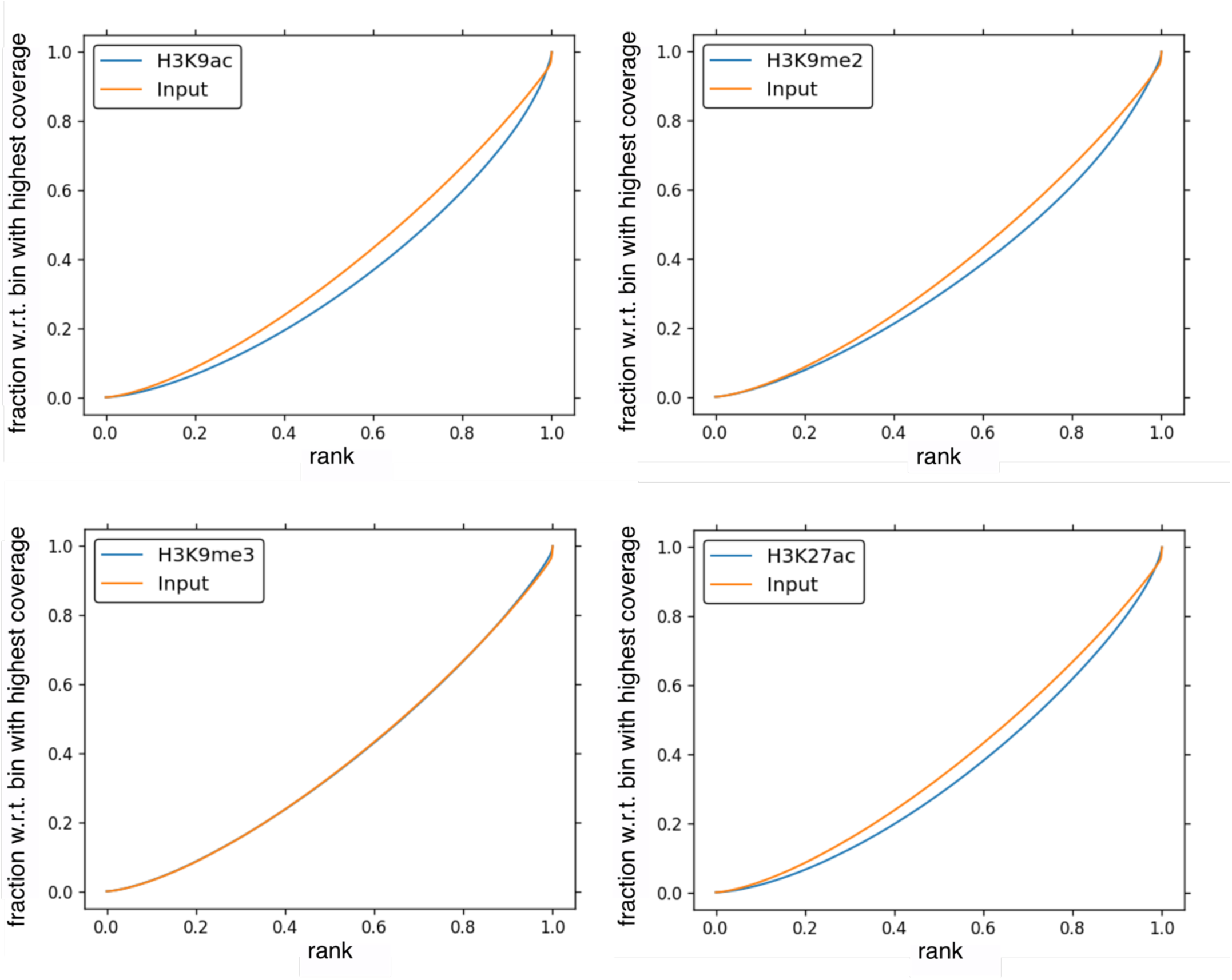
Enrichment strength of ChIP signals in 9-WT input and histone modifications.

**Figure S11.**
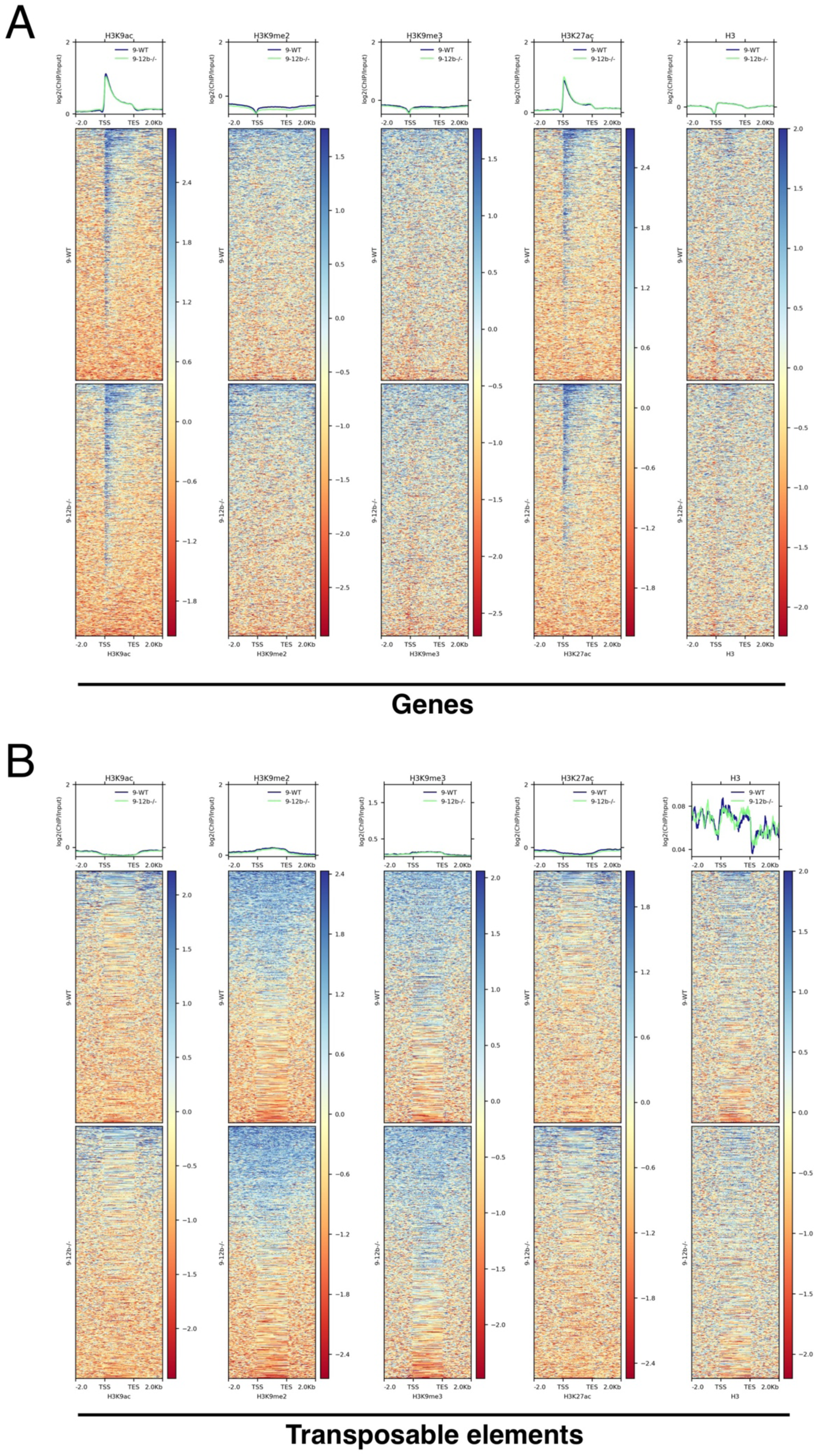
Heatmaps of genome-wide histone modification enrichments in genes and transposable elements (TEs) of rice 9-WT and *hac701* (*9-12b^-/-^*) mutant. Enrichment of H3K9 acetylation, H3K9 di-methylation, H3K9 tri-methylation, H3K27 acetylation, and H3 in genes (*A*) and TE (*B*) regions. Data represent one of the two biologically independent samples showing similar enrichment results.

**Figure S12.**
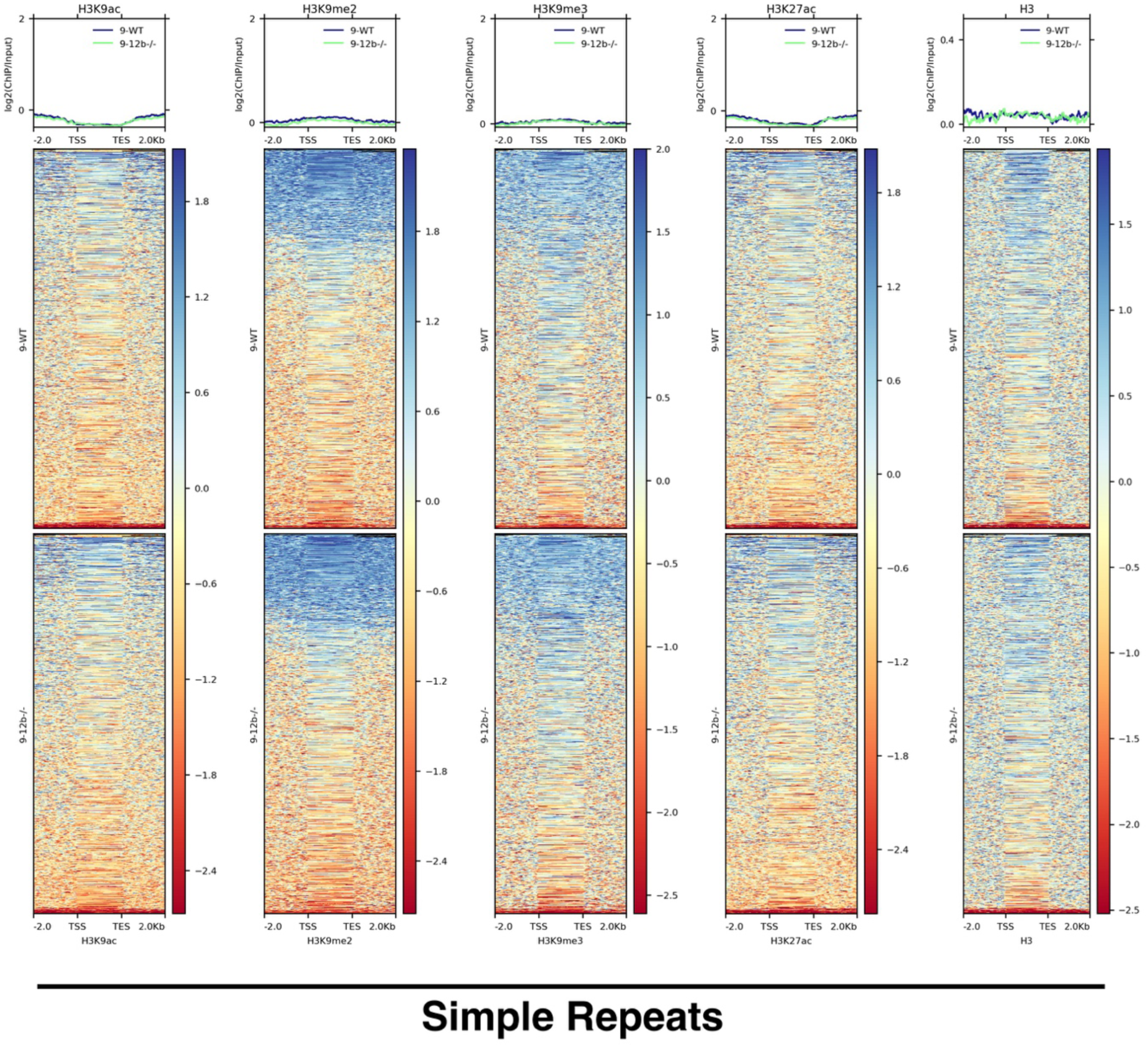
Heatmaps of genome-wide histone modification enrichments in simple repeats of rice 9-WT and *hac701* (*9-12b^-/-^*) mutant. Enrichment of H3K9 acetylation, H3K9 di-methylation, H3K9 tri-methylation, H3K27 acetylation, and H3 in simple repeat regions. Data represent one of the two biologically independent samples showing similar enrichment results.

**Figure S13.**
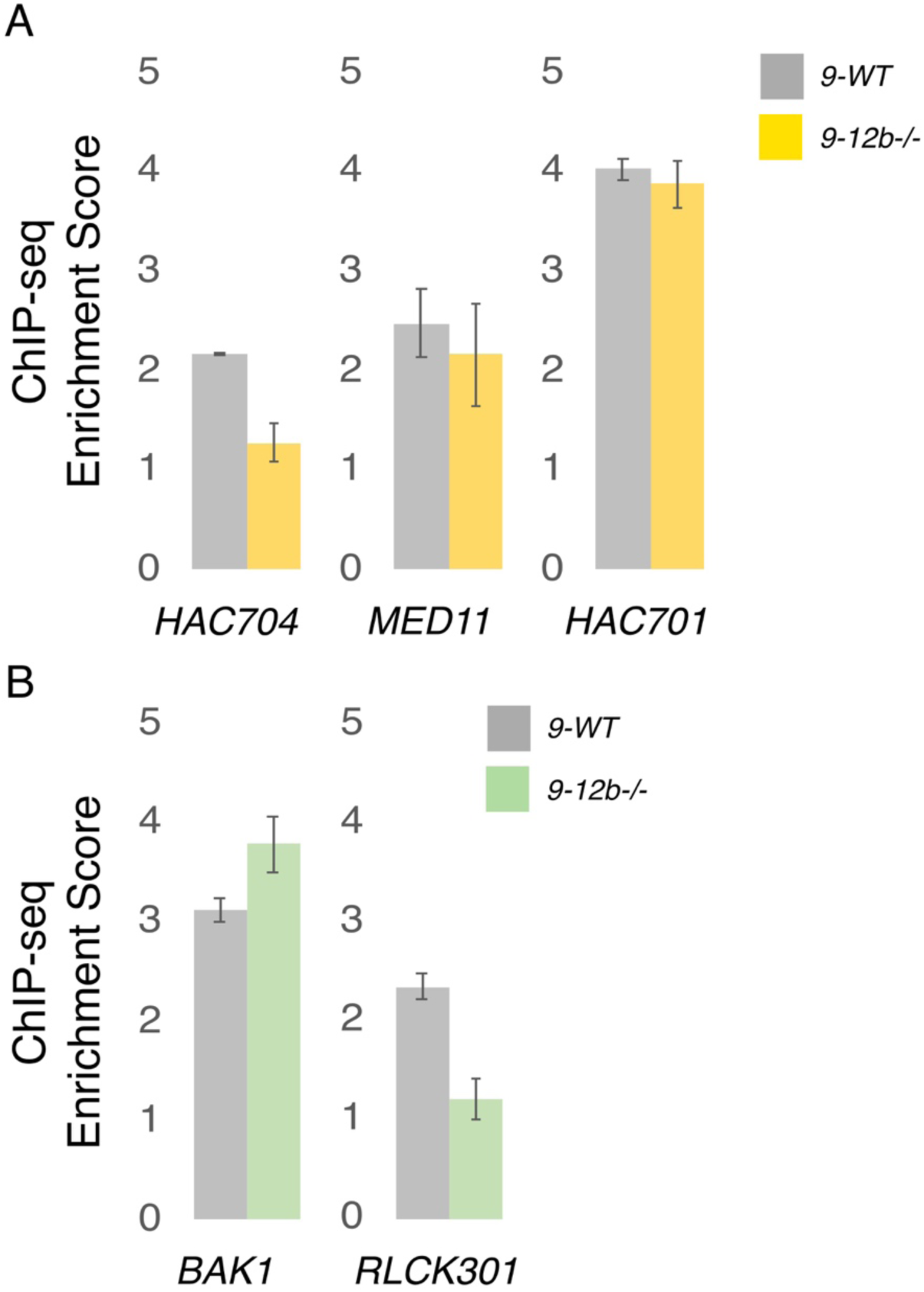
Enrichment scores of additional representative genes showing differential enrichments of H3K9 acetylation in *hac701* (*9-12b^-/-^*) mutant. (*A*) Transcriptional coactivator genes showing reduced or partially reduced H3K9 acetylation enrichments. (*B*) Putative guardee genes showing differential enrichments of H3K9 acetylation. Scores are averages of two biologically independent ChIP-seq replicates with bars showing standard deviation (SD).

**Figure S14.**
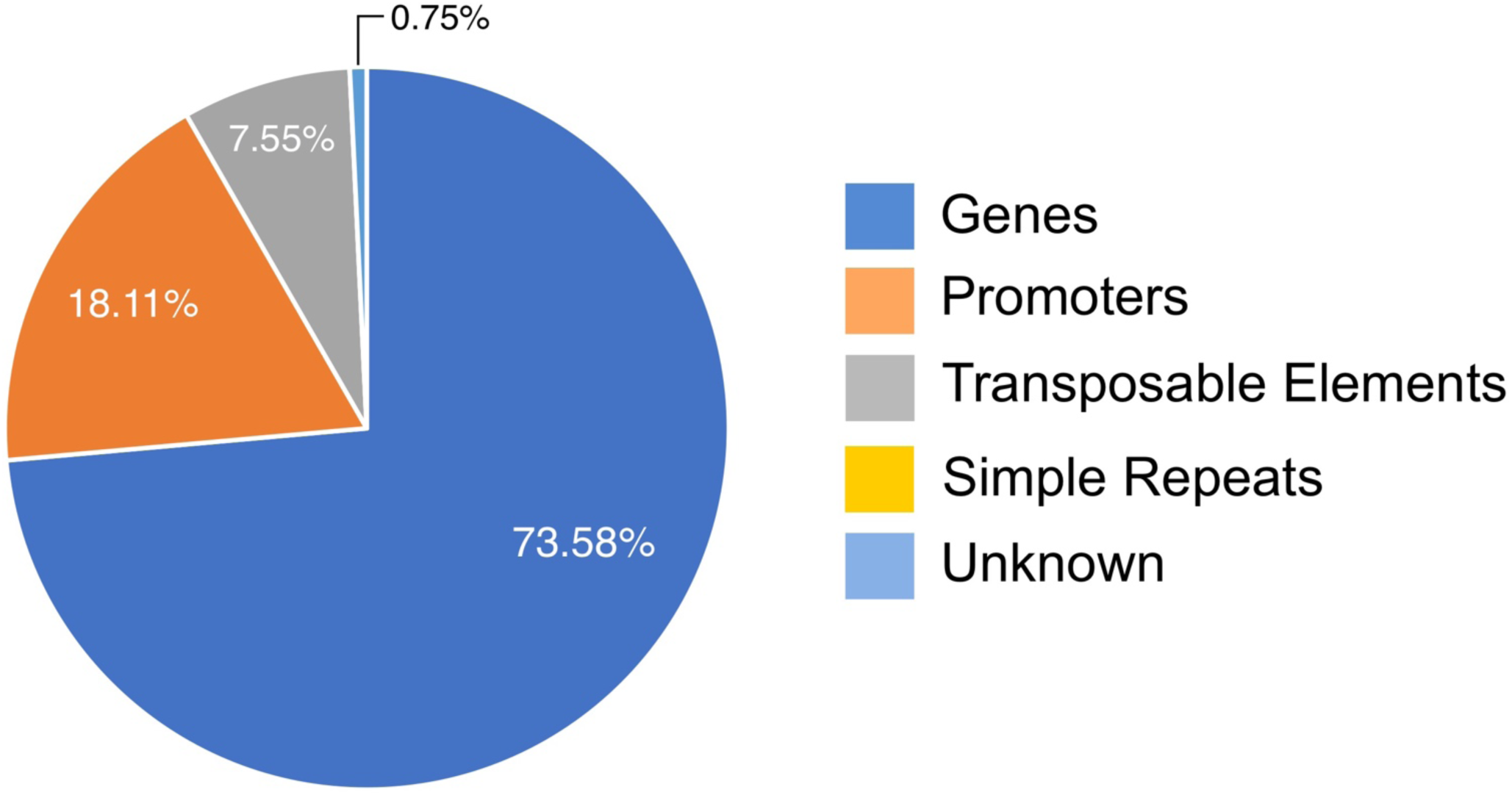
Distribution on genomic locations of 265 regions in H3K9 acetylation ChIP (wild type/mutant).

**Figure S15.**
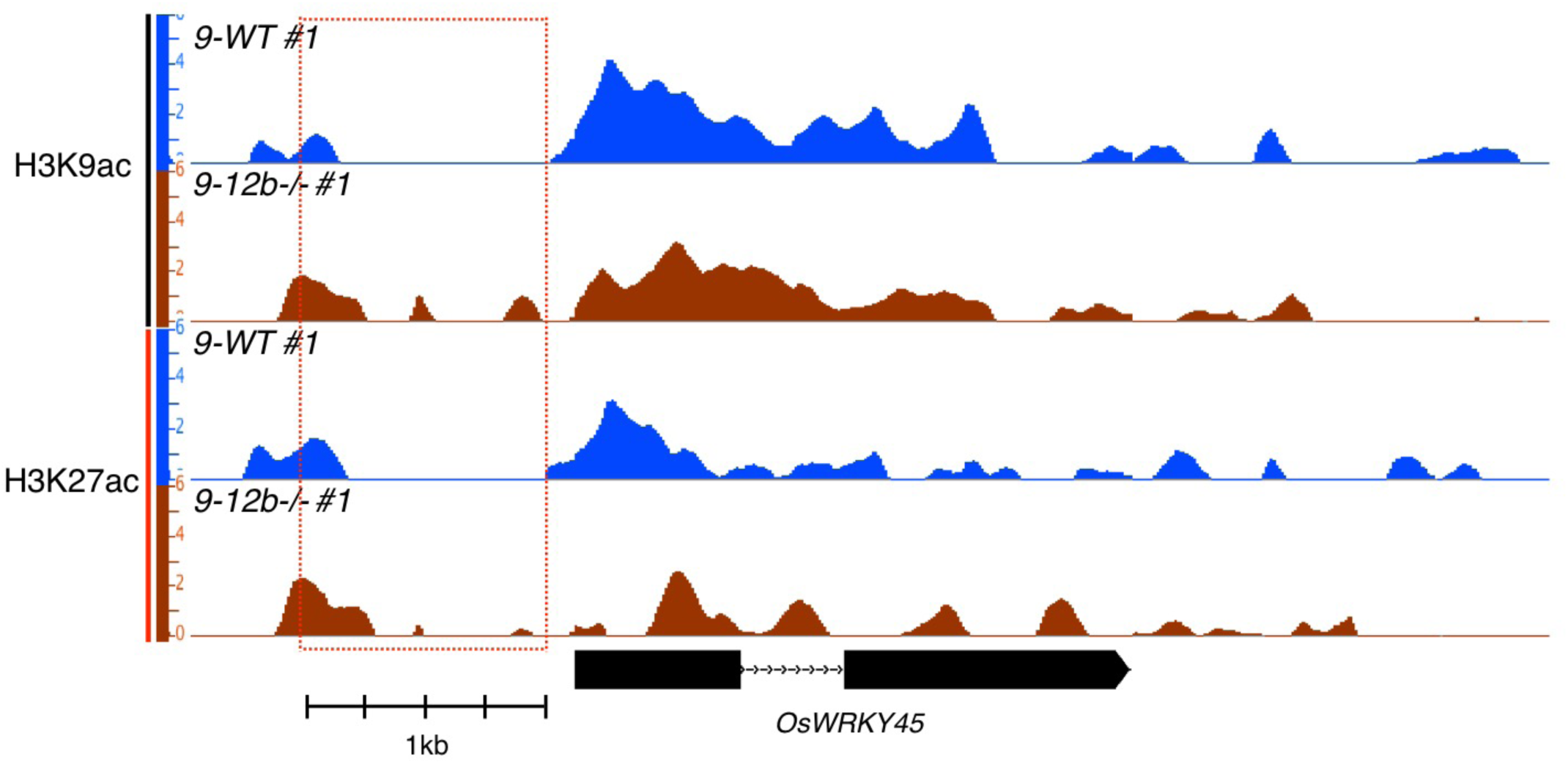
O*s*WRKY45 depth graph of H3K9 and H3K27 acetylation ChIP-seq showing the number of reads in RPKM (Reads Per Kilobase of transcript, per Million mapped reads) in 9-WT and *hac701* (*9-12b^-/-^*) mutant. The red dotted box shows the 1 kb upstream region of *OsWRKY45* with arrow indicating the direction of transcription. Data show a representative sample.

**Figure S16.**
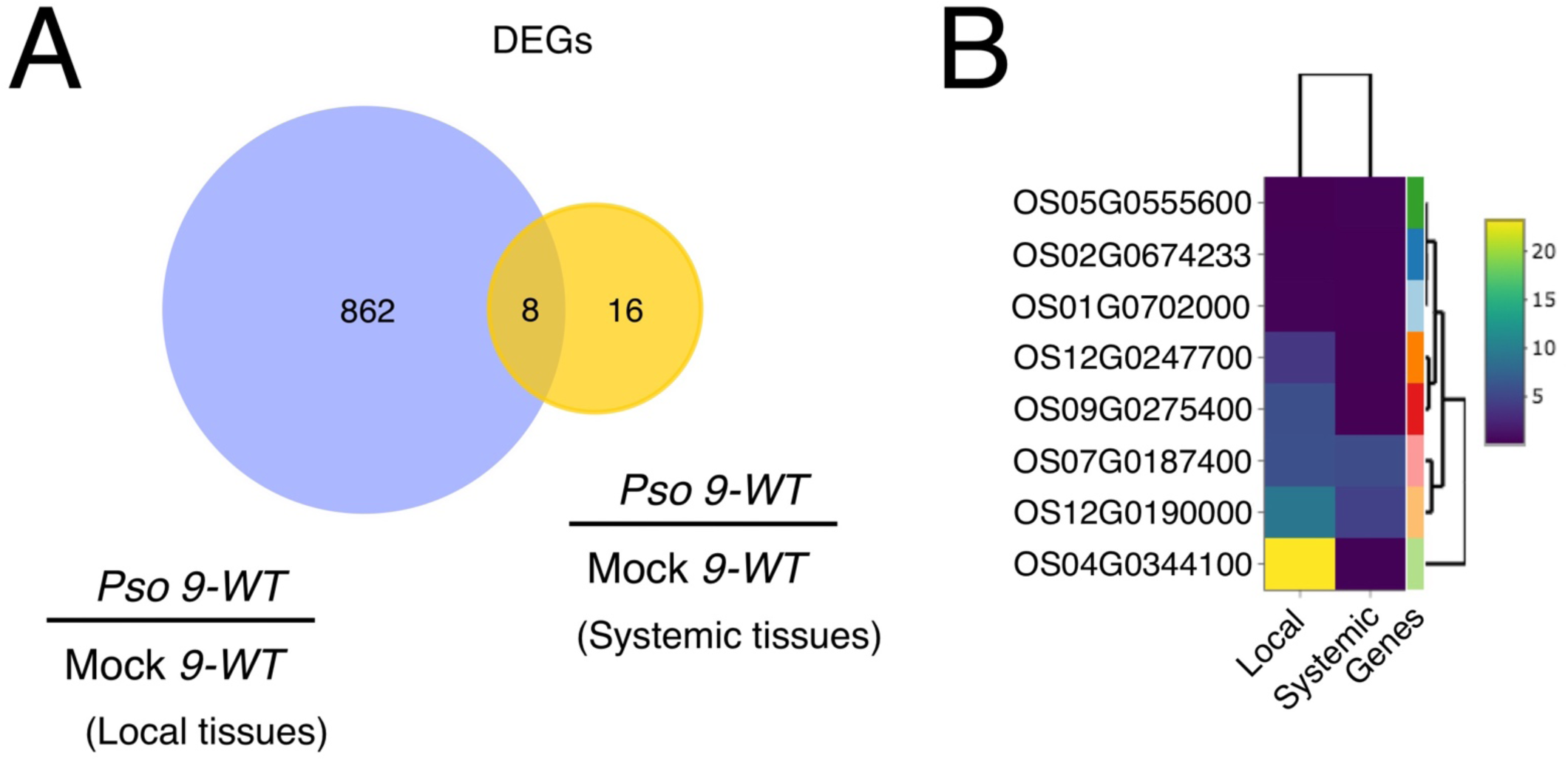
Systemic gene expression analysis in rice-*Pseudomonas syringae* pv. *oryzae* (*Pso*) pathosystem. (*A*) Differentially expressed genes (DEGs) in the local (870) and systemic (24) tissues of plants infected with *Pso* (Table S16). Eight DEGs were found to be common in both local and systemic tissues. (*B*) The expression of eight DEGs shown in log2 fold changes. For local tissue analysis, two independent RNA-seq samples in both treatments (mock and *Pso*) are presented. For systemic tissue analysis, two independent RNA-seq samples in mock and three in *Pso* treatment are presented. The number of reads are in RPKM (Reads Per Kilobase of transcript, per Million mapped reads) in 9-WT and *hac701*.

**Figure S17.**
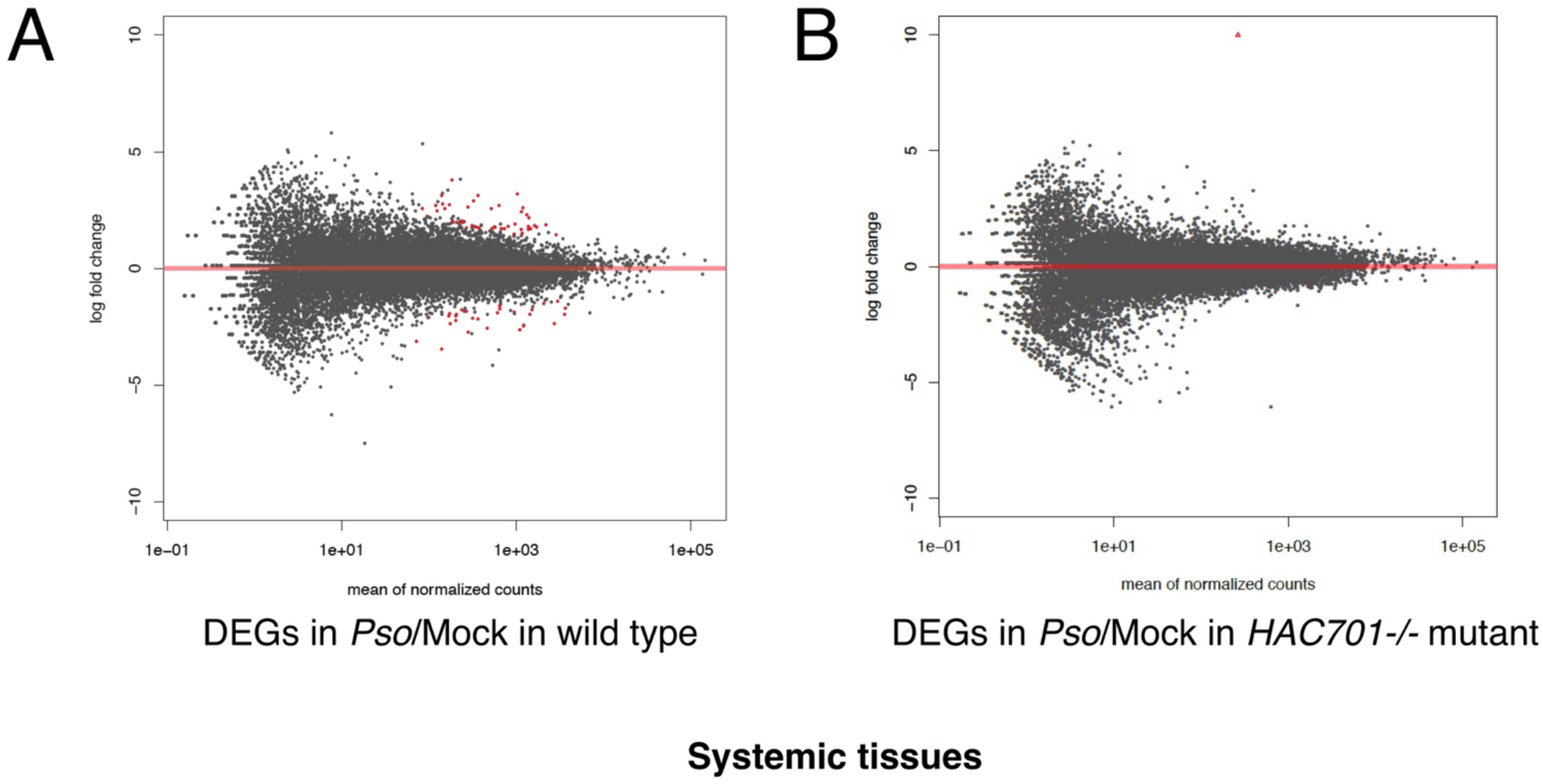
MA-plot of differentially expressed genes (DEGs) in systemic tissues of 9-WT and *hac701* infected with *Pseudomonas syringae* pv. *oryzae* (*Pso*). (*A*) and (*B*) plots showing differences in measurements of differentially expressed genes (DEGs). *P* adjusted value is set at <0.01. For 9-WT and *hac701* samples, two independent RNA-seq samples in mock treatment and three in *Pso* treatment are presented.

**Figure S18.**
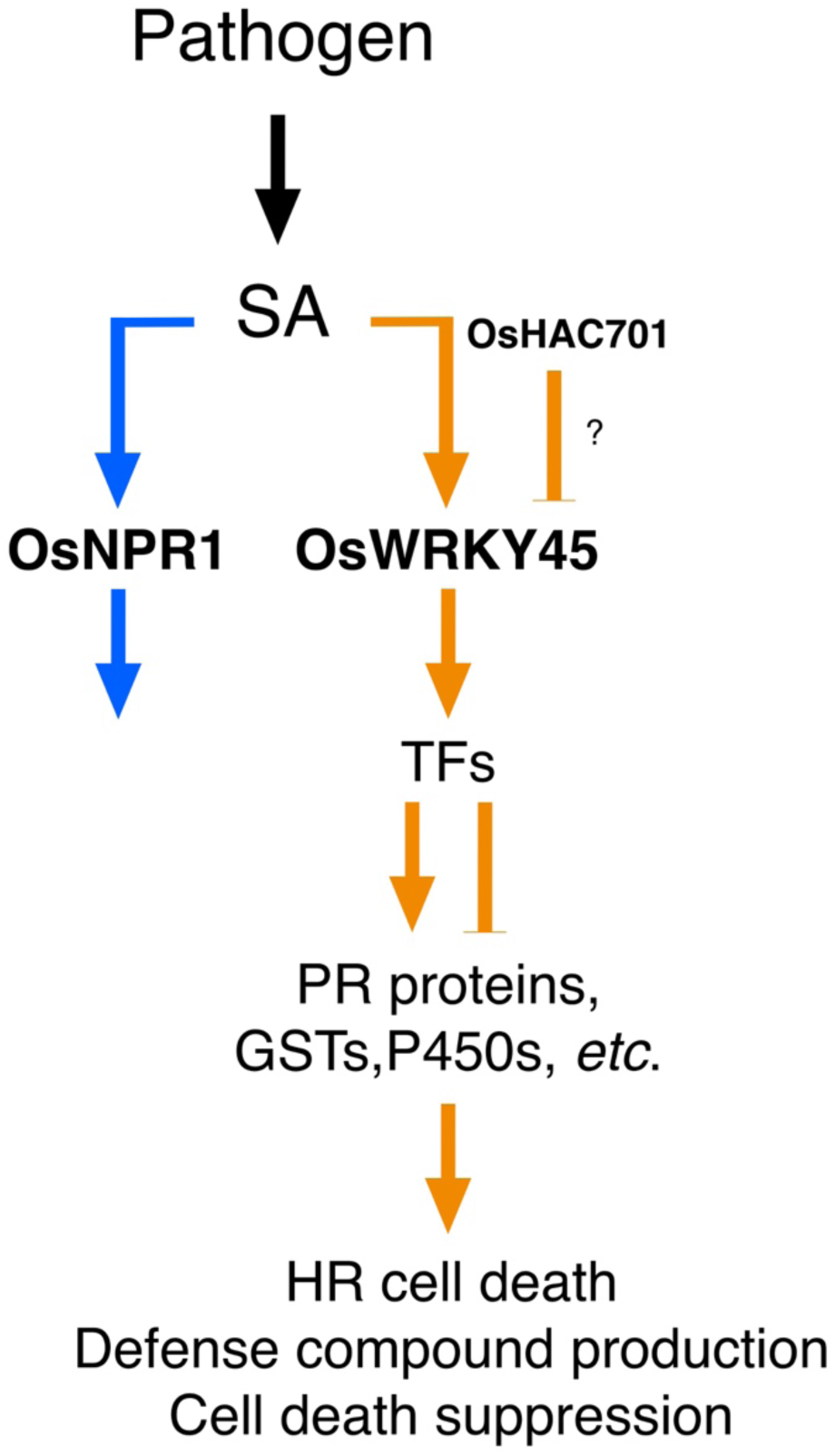
O*s*WRKY45-dependent defense pathway is suppressed by *HAC701*. Defense-related genes regulated directly and indirectly by *OsWRKY45* are negatively regulated by *HAC701* resulting in resistance phenotype of *hac701* through *OsWRKY45* upregulation.

## SUPPLEMENTARY TABLE LEGENDS

Table S1 sgRNA oligonucleotides used for CRISPR/Cas9 mutagenesis and primers for genotyping. Bold sequences in the guide RNAs are the Protospacer Adjacent Motif (PAM).

Table S2 Summary of total RNA-sequencing data in local tissues.

Table S3 Wild type (9-WT) differentially expressed genes (DEGs) in *Pso*.

Table S4 *HAC701* repressed and enhanced genes.

Table S5 HAC701 mutation differentially expressed genes (DEGs) in Mock (*HAC701mut*-DEGs in Mock).

Table S6 *HAC701* mutation differentially expressed genes (DEGs) in *Pso* (*HAC701mut*-DEGs in *Pso*). The motifs were identified 1kb upstream sequence of the start codon.

Table S7 BTH-responsive genes (24 hpi) in microarray experiments (34, 35) (Shimono et al. 2007, The Plant Cell & Sugano et al. 2010, Plant Molecular Biology). Overlap genes were removed.

Table S8 BTH-inducible & WRKY45-dependent genes (33) (Nakayama et al. 2013, BMC Plant Biology).

Table S9 Upregulated genes after DEX-induced *WRKY45* expression (33) (Nakayama et al. 2013, BMC Plant Biology).

Table S10 BTH-inducible genes affected by rice *NPR1/NH1* knockdown (35) (Sugano et al. 2010, Plant Molecular Biology).

Table S11 HAC701 mutation differentially expressed genes (DEGs) found in BTH-WRKY45 regulated genes that contain W-box motifs. The motifs were identified 1 kb upstream sequence of the start codon.

Table S12 *HAC701* mutation differentially expressed genes (DEGs) found in BTH-WRKY45 and BTH-NPR1/NH1 regulated genes that contain W-box motifs. The motifs were identified 1 kb upstream sequence of the start codon.

Table S13 Significantly enriched DNA motifs in HAC701 mutation differentially expressed genes (DEGs) in *Pso* (HAC701mut-DEGs in *Pso*).

Table S14 Rice ChIP-sequencing information of histone modifications.

Table S15 Identified peaks in H3K9ac and their corresponding genes in 9-WT/*9-12b^-/-^* (wild type/mutant). Colored boxes indicate the repeated genes.

Table S16 Summary of total RNA-sequencing data in systemic tissues.

Table S17 List of R genes upregulated in *hac701* in *Pso*.

Table S18 Identified peaks in H3K9me3 and their corresponding genes in *9-12b^-/-^*/Input (mutant/Input). Colored boxes indicate repeated genes.

Table S19 Oligonucleotides used for RT-qPCRs (1) and ChIP-qPCRs (2).

## ACKNOWLEDGEMENTS

We thank OIST SQC for RNA-seq and ChIP-seq analysis. This project benefited greatly from the comments of Drs. Yusuke Saijo, Hirofumi Nakagami, Justin Walley, Yasuhiro Kadota, and from the members of the Plant Epigenetics Unit (OIST) and Plant Immunity Research Group (RIKEN CSRS).

## FUNDING

This work was supported in part by the Okinawa Institute of Science and Technology Graduate University (OIST). N.A.E. was also funded by Grant-in-Aid for JSPS Fellows (Grant No: 18F18085). The funders had no role in study design, data collection and analysis, decision to publish, or preparation of the manuscript.

## AUTHOR CONTRIBUTIONS

NAE: Conceptualization, Data curation, Funding acquisition, Formal analysis, Investigation, Visualization, Writing-original draft, Writing-review & editing

LNT: Data curation, Formal analysis, Writing-review & editing

SM: Formal analysis, Investigation

YS: Formal analysis, Investigation

KS: Funding acquisition, Formal analysis, Supervision, Writing-review & editing

HS: Funding acquisition, Formal analysis, Investigation, Supervision, Writing-review & editing

## COMPETING INTERESTS

The authors have declared that no competing interests exist.

## Notes

https://osf.io/4un59/

### Competing Interest Statement

The authors have declared no competing interest.

